# Relaxase Asymmetry Drives Initiation of Bacterial Conjugation

**DOI:** 10.64898/2026.02.20.706640

**Authors:** Danylo Gorenkin, Ruina Liang, Jonasz B. Patkowski, Salomé Bodet-Lefèvre, Tiago R. D. Costa, Aravindan Ilangovan

## Abstract

Bacterial conjugation is the major route of horizontal gene transfer and a key driver of the dissemination of antibiotic resistance, exacerbating the global health crisis. Central to this process is TraI, a bifunctional relaxase-helicase encoded by the iconic F-plasmid family. TraI initiates single-stranded DNA (ssDNA) transfer by covalently attaching to plasmid DNA through its trans-esterase (TE) activity and subsequently unwinds the plasmid via its helicase activity. Although genetic and biochemical studies have postulated that two TraI molecules are required for efficient conjugative transfer, how these molecules coordinate their activities in space and time - and how strand nicking is coupled to helicase loading - has remained unknown.

Here, we capture the elusive, functionally asymmetric TraI homodimer and report its cryo-EM structure, revealing that the interaction between the two TraI molecules is mediated predominantly by ssDNA. We further show that TE-driven duplex melting at the origin of transfer (*oriT*) by one TraI molecule enables helicase loading by a second TraI molecule on the transfer strand of the unnicked plasmid DNA. Although the TE domain is intrinsically competent for strand nicking, its activity is inhibited by the host factor IHF, and this inhibition is relieved upon helicase loading. Together, these findings define a regulatory loop that coordinates helicase loading with strand nicking, providing mechanistic insight into the earliest stages of conjugative DNA processing.

## Introduction

Antibiotic resistance is a mounting global health crisis, with drug-resistant infections now reported across virtually all major pathogen classes and steadily eroding the efficacy of both first-line and last-resort antibiotics^1^. A principal mechanism driving this spread is bacterial conjugation, which enables horizontal transfer of antibiotic-resistance determinants carried on plasmids or on genomic elements from a donor to a recipient cell^2–4^. First discovered in the 1940s, conjugation not only revealed DNA exchange between bacteria but also seeded many of the foundational genetic experiments in *Escherichia coli* that shaped modern molecular biology^5^. Mechanistically, conjugation is orchestrated by three tightly coordinated modules: (i) a multi-megadalton membrane-embedded type IV secretion system (T4SS)^6^; (ii) an extended conjugative pilus, built from a complex of pilin and phospholipid^7^, that physically bridges donor and recipient bacterial cells^8^; and (iii) the relaxosome, a nucleoprotein complex that processes plasmid DNA to prepare it for transfer^5,9^. Over the past decade, structural and biochemical studies have illuminated key aspects of T4SS and pilus assembly, offering a detailed understanding of how these nanomachines couple cells and translocate macromolecular cargo. In contrast, the molecular events underlying relaxosome-mediated DNA processing – particularly the order, regulation, and physical coordination of enzymatic steps at the origin of transfer (*oriT*) – remain less understood^5,10^. Recent cryo-EM analyses provided the first structural snapshots of the F-plasmid relaxosome, revealing the interaction networks that stabilize the complex^11^. The relaxosome of the most widely studied F/R1 plasmid family, comprises plasmid-encoded accessory proteins TraM and TraY, together with the host-encoded architectural factor IHF. These proteins recruit the main effector the relaxase (TraI) to its cognate binding site on *oriT*, initiating DNA processing^9^.

Relaxases are central to conjugation and are critical for DNA processing and subsequent transfer, possibly piloting the transfer strand (T-strand) to the recipient cell. In the F/R1 plasmid system, the relaxase (TraI) is a large (~200 kDa) protein with an N-terminal trans-esterase (TE) domain and two central helicase domains belonging to the SF1B helicase subfamily (Figure 1A). The TE domain recognizes the ‘*TE-binding site*’ within *oriT*, located immediately upstream (5′) of the nick site (*nic*), where it cleaves the T-strand and forms a covalent phosphotyrosyl linkage via an active-site tyrosine. Meanwhile, the helicase domains engage the region immediately downstream (3′) of *nic*, termed the ‘*helicase-binding region*’ (Figure 1C). Once loaded, TraI functions as a monomeric helicase, unwinding plasmid DNA to generate the T-strand ssDNA substrate for translocation to the recipient cell through the type IV secretion system (T4SS). Critically, the TE and helicase activities of TraI are mutually exclusive within a single molecule – a single TraI protomer cannot simultaneously engage both the *TE-binding site* and *helicase-binding regions*^12,13^. Therefore, two separate TraI molecules must act in concert at *oriT*, each assigned to one of the adjacent binding sites. This model is supported by biochemical data showing a dimeric TraI intermediate on a ssDNA substrate spanning the *TE-binding site*, *nic*, and the *helicase-binding region.* The TE-engaged TraI adopts an open conformation, while the helicase-engaged protomer assumes a closed, helicase-active conformation^13^. Together, this configuration forms a ‘functionally asymmetric homodimer’ during initiation of conjugation. This asymmetry during initiation contributes towards the eventual dual roles of TraI in the donor and recipient cells: one protomer is transferred with the T-strand into the recipient, where it facilitates re-ligation and circularization of the plasmid, while the second protomer remains in the donor to process the replacement strand.^10,14^. While the requirement for two TraI molecules is now well accepted, the molecular basis of their coordination – how they assemble at *oriT*, divide labour, and temporally regulate strand processing – remains poorly defined. The structural architecture of this functionally asymmetric dimer and the regulatory mechanisms that gate initiation of conjugation are still unclear.

**Figure 1.**
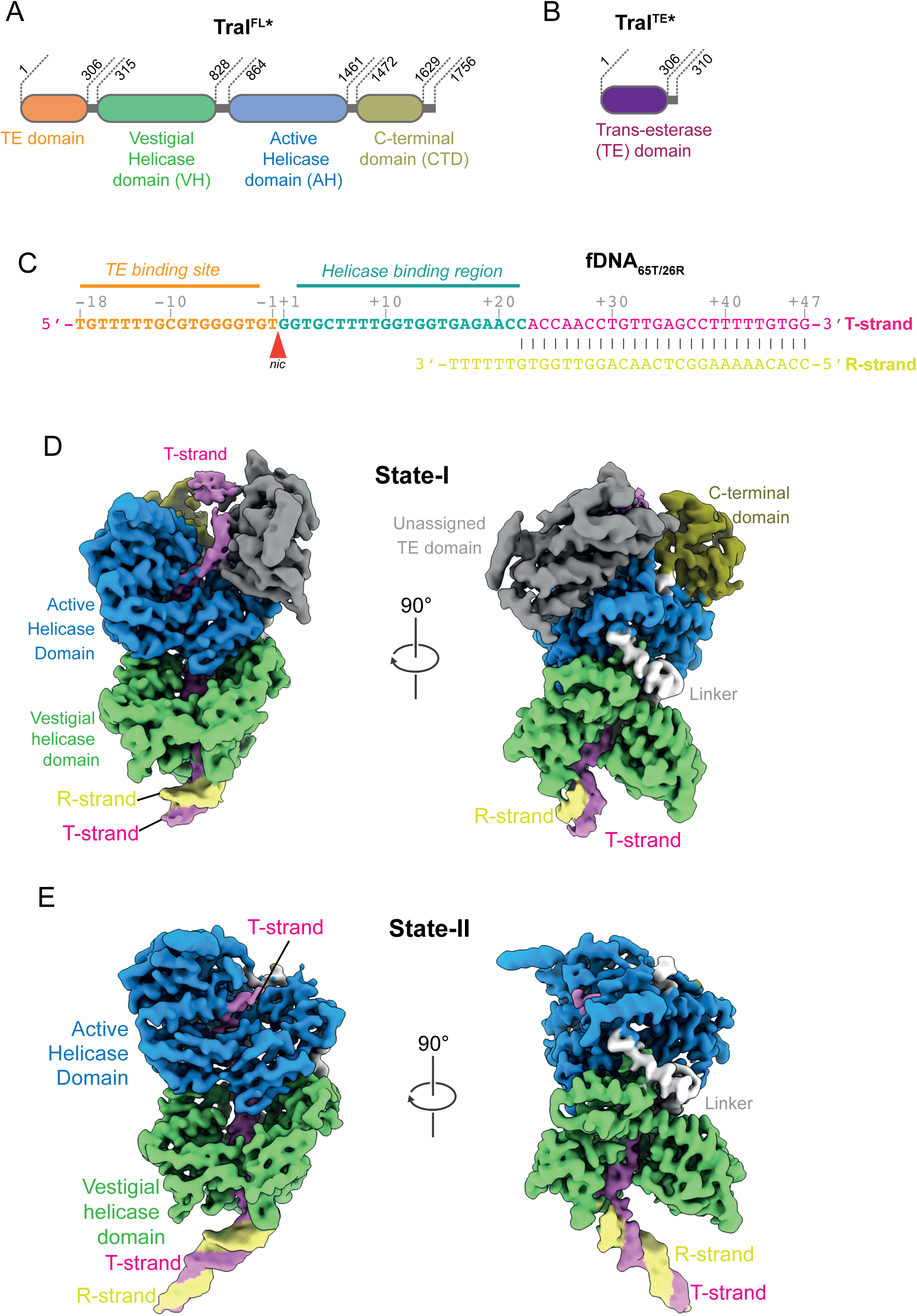
Biochemical reconstitution and cryo-EM structures of the TraI^TE*^–TraI^FL*^–fDNA65T/26R complex. a. Domain organisation of the TraI^FL*^ construct used for biochemical assembly, with domain boundaries. b. Domain organisation of the TraI^TE*^ construct used for biochemical assembly, highlighting individual domains and their boundaries. c. Sequence of the DNA fork substrate (fDNA65T/26R) used for assembly of the tripartite TraI^TE*^:TraI^FL*^:fDNA65T/26R complex. d. Cryo-EM density map corresponding to class-1, showing the active helicase (AH) domain (blue), vestigial helicase (VH) domain (green), C-terminal domain (CTD; olive green), helicase linker (light grey), DNA duplex (R-strand, yellow; T-strand, pink). The TE domain is not assigned in this reconstruction and is shown in dark grey. Two views are presented, rotated 90° relative to each other. e. Cryo-EM density map corresponding to class-2, showing the AH (blue) and VH (green) domains, helicase linker (light grey), DNA duplex (R-strand, yellow; T-strand, pink), and T-strand ssDNA (pink). Two views are shown, rotated by 90°.

Here, we define these mechanisms by solving high-resolution cryo-EM structures of TraI bound to a forked DNA substrate that emulates the native *oriT’s* architecture. These structures resolve the spatial arrangement of the TE and helicase modules across a duplex– ssDNA junction and define how the TE domain from one molecule of TraI accesses the *oriT* 5′ of *nic*, while the helicase domain from a second molecule of TraI binds to the region 3′ of *nic*. Using supercoiled plasmid DNA together with rationally designed oligonucleotide substrates, we further define how *oriT* recruits two TraI protomers and how each adopts either a TE-engaged or helicase-engaged configuration. We identify features of the relaxosome that stabilize a pre-activation state and demonstrate that helicase loading precedes nicking, coordinated by the host factor, IHF. Collectively, our findings reveal how functional asymmetry is established within a TraI dimer, how enzymatic activities are temporally partitioned, and how relaxosome-mediated regulation primes *oriT* for conjugative transfer. This work provides a mechanistic framework for understanding the earliest events in bacterial conjugation and offers new insight into how DNA-processing enzymes can be regulated through spatial segregation and host-factor gating. Given the central role of conjugation in the horizontal spread of antibiotic resistance, these findings advance our understanding of a fundamental biological process.

## Results

### Biochemistry and cryo-EM structure determination

Two molecules of TraI participate in conjugative DNA transfer and come together at the initial step of DNA processing as a functionally asymmetric homodimer^12,13^. To understand the molecular architecture of this asymmetric relaxase dimer-DNA complex, we reconstituted a minimal functional assembly in which one TraI molecule is represented by the full-length TraI (TraI^FL^; Figure 1A), while the second molecule is the isolated TE domain construct (TraI^TE^; Figure 1B). We have used the catalytically inactive Y16F/Y17F mutant in both these constructs, which henceforth will be denoted as TraI^TE*^ and TraI^FL*^. As the substrate for complex formation, we employed a DNA fork in which the T-strand included the *TE binding site*, the *nic site* and the *helicase binding region*, presented in single-stranded form, with an additional 25 base pairs after the 3’ end of the *helicase binding region* (65 bases). The replicative (R-strand) includes nucleotides complementary to the 25 bases outside the *helicase binding region* with a 3’ 6 base poly-T tail (34 bases). The T- and R-strands are maintained as duplex DNA, henceforth called fDNA65T/26R (Figure 1C). The tripartite (TraI^TE*^–TraI^FL*^–fDNA65T/26R) assembly was reconstituted *in vitro* and purified by size-exclusion chromatography, yielding two distinct nucleoprotein complexes corresponding to a higher molecular weight TraI^TE*^:TraI^FL*^:fDNA65T/26R complex and a lower molecular weight TraI^TE*^:DNA complex, respectively (Figure S1A). The TraI^TE*^:TraI^FL*^:fDNA65T/26R complex corresponding to the first peak was used for cryo grid preparation, single-particle cryo-EM data collection, and image processing (Figure S1A, S1B, S1C and S2). During data processing, rounds of 3D classification combined with heterogenous refinement produced two distinct reconstructions (Figure S2). The first class (class 1) shows clear density for both duplex DNA and the TE domain, consistent with TraI adopting the helicase engaged conformation observed previously and we henceforth refer to this as state-1^13^ (Figure 1D and S2). The second reconstruction, comprising classes 2 and 3, is largely similar to the class 1, but lacks density attributable to the characteristic TE domain, and we henceforth refer to this as state-2 (Figure 1E and S2).

### State-1 is a single TraI molecule in its helicase mode

State-1 displays a single TE domain as opposed to the expected two TE domains, suggested by the biochemical reconstitution experiments (Figure 1D, S1A and S1B). To identify the TraI molecule that contributed the TE domain, i.e. whether the TE is provided *in trans* by a separate molecule (TraI^TE*^), or *in cis* by the full-length molecule (TraI^FL*^), we examined the density map while systematically increasing the sigma threshold from 3.0σ to 1.0σ. As the threshold was increased, additional density appeared and eventually connected with the vestigial helicase (VH) domain (green), thereby establishing a direct link between the helicase core and the unidentified TE density (grey) (Figure S5A and S5B). This analysis confirmed that the TE domain observed in state-1 is part of the full-length TraI construct (TraI^FL*^) used to reconstitute the complex, which adopts the helicase mode and will be referred as TraIHEL (Figure 2A and 2B). To interpret the interaction between fDNA65T/26R and the helicase core of TraI, we used the high-resolution map obtained from local refinement, focused on the helicase domains without the TE domain using 1.3 million particles (TraIHEL-FOC; Figure S2). This map corresponds to the combined reconstruction prior to the classification of particles into state-1 and state-2. Owing to the higher resolution at the helicase core, the purine and pyrimidine rings are well resolved, allowing unambiguous assignment of the ssDNA sequence bound within the helicase core and extending through the entire fDNA65T/26R template used. The DNA trajectory in this map shows a marginal deviation compared to previous reconstructions, including a small bend primarily introduced by T12, which was not visible before^13^. Additionally, G11 adopts two alternate conformations, both of which are clearly supported by the density (Figure 2C). This structure captures TraI in the helicase mode bound to a DNA fork substrate, closely mimicking the native template during conjugation. Residues R664, K723 and R728 make direct contacts with the double-stranded region of the DNA, whereas W659 lies closer to the duplex-single-strand junction where the strands separate (Figure 2D). To investigate the role of specific residues in the TraI-DNA interaction, we performed a series of *in vivo* conjugation assays using different TraI point mutants (Figure S5E and S5F). None of the tested point mutations exhibited a statistically significant difference in conjugation efficiency compared to the wild-type TraI, consistent with our structure showing that TraI and DNA interact across the majority of the *helicase binding region.* This finding implies that a more extensive series of mutations would be necessary to fully disrupt TraI’s overall DNA binding and helicase function. This is exemplified by the results of conjugation efficiency with R1520A and K1521A single mutants, which show native conjugation efficiency levels compared to a R1520A+K1521A double mutant, which showed a modest decrease of one order of magnitude in efficiency (Figure S5E and S5F).

**Figure 2.**
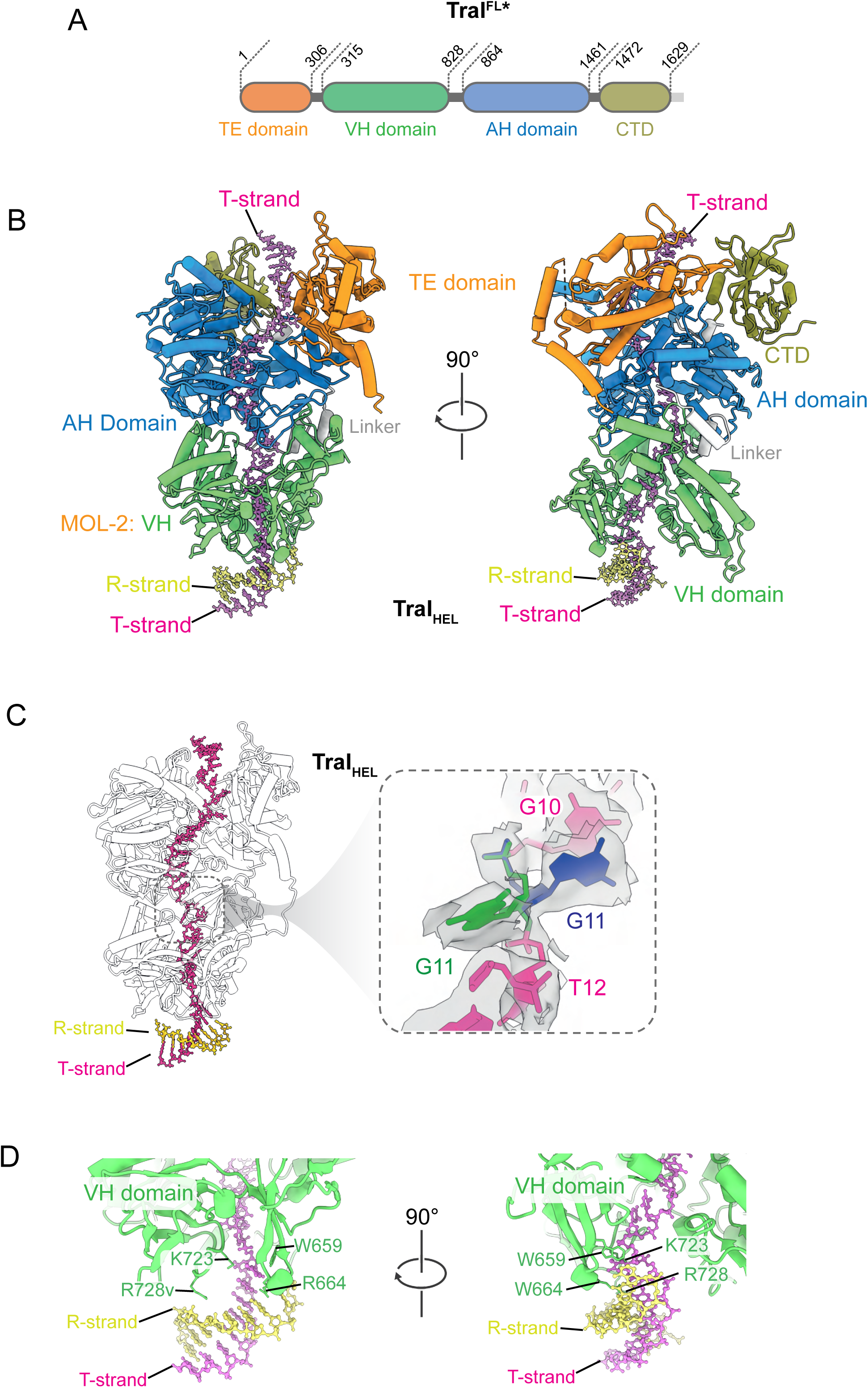
Class-1 represents TraI in the helicase-loading mode. a. Domain organisation of the TraI^FL*^ construct used in the biochemical assembly, with boundaries indicated. Sequence C-terminal to the CTD (beyond residue 1629) is not resolved in the cryo-EM map and shown in light grey. b. Cryo-EM structure of the TraIHEL complex, showing TraI^FL*^ bound to the forked DNA substrate (fDNA65T/26R). Domains are coloured as follows: TE (orange), vestigial helicase (VH; green), active helicase (AH; blue), and CTD (olive green). Two views are shown, rotated by 90°. c. TraIHEL structure with the protein rendered in grey and the DNA highlighted. The T-strand and R-strand of the forked DNA are coloured yellow and pink, respectively. Inset: density revealing dual occupancy at nucleotide position G11. d. Interaction between the DNA duplex and the vestigial helicase domain mediated by residues W659, R664, R728 and K723.

Second, density corresponding to the previously unresolved portion of the C-terminal domain (CTD, residues 1472–1628) is now visible, although the extreme C-terminal residues (1629–1756) remain disordered (Figure 2A and 2B). Upon closer inspection, we observed an additional density positioned between the CTD and the TE domain. As the map threshold was increased (σ = 0.25), this density became continuous, suggesting it represents a feature of the structure (Figure S5C). When the atomic model is overlaid, it appears that this density is consistent with the path of the ssDNA exiting the active helicase domain after passing near the TE domain as observed before^13^ (Figure S5D). As this structure of TraIHEL solely contains one molecule of TraI^FL*^ bound to the forked DNA substrate, the ssDNA binding site of the TE domain remains unoccupied. We therefore propose that the additional density represents the ssDNA exiting the active helicase domain and appears to also transiently interact with the CTD, potentially involving residues R1520 and K1521 (Figure S5D). Given the low-resolution nature of the map at this region, it is however not possible to resolve interactions at the specific residue/base level.

### State-2 is the asymmetric homodimer, the dimer intermediate of TraI

At a stringent contour level (σ = 0.4), where high-resolution features can be individually resolved, the map of state-2 shows no discernible density for a TE domain (Figure 1E). As the contour is lowered, density emerges that extends from the exiting ssDNA to the active helicase domain and grows into a larger globular feature. By σ = 0.14 this density forms a continuous connection to the main TraI-helicase density via the ssDNA path (Figure 3A). Although this extra mass linked to the helicase by ssDNA could represent a TE domain, only a single globular feature is observed even though two TE domains are present in the biochemical assembly (Figure S1A and S1B). Due to poor local resolution in this region of the map an unambiguous assignment based solely on contour manipulation is not possible.

**Figure 3.**
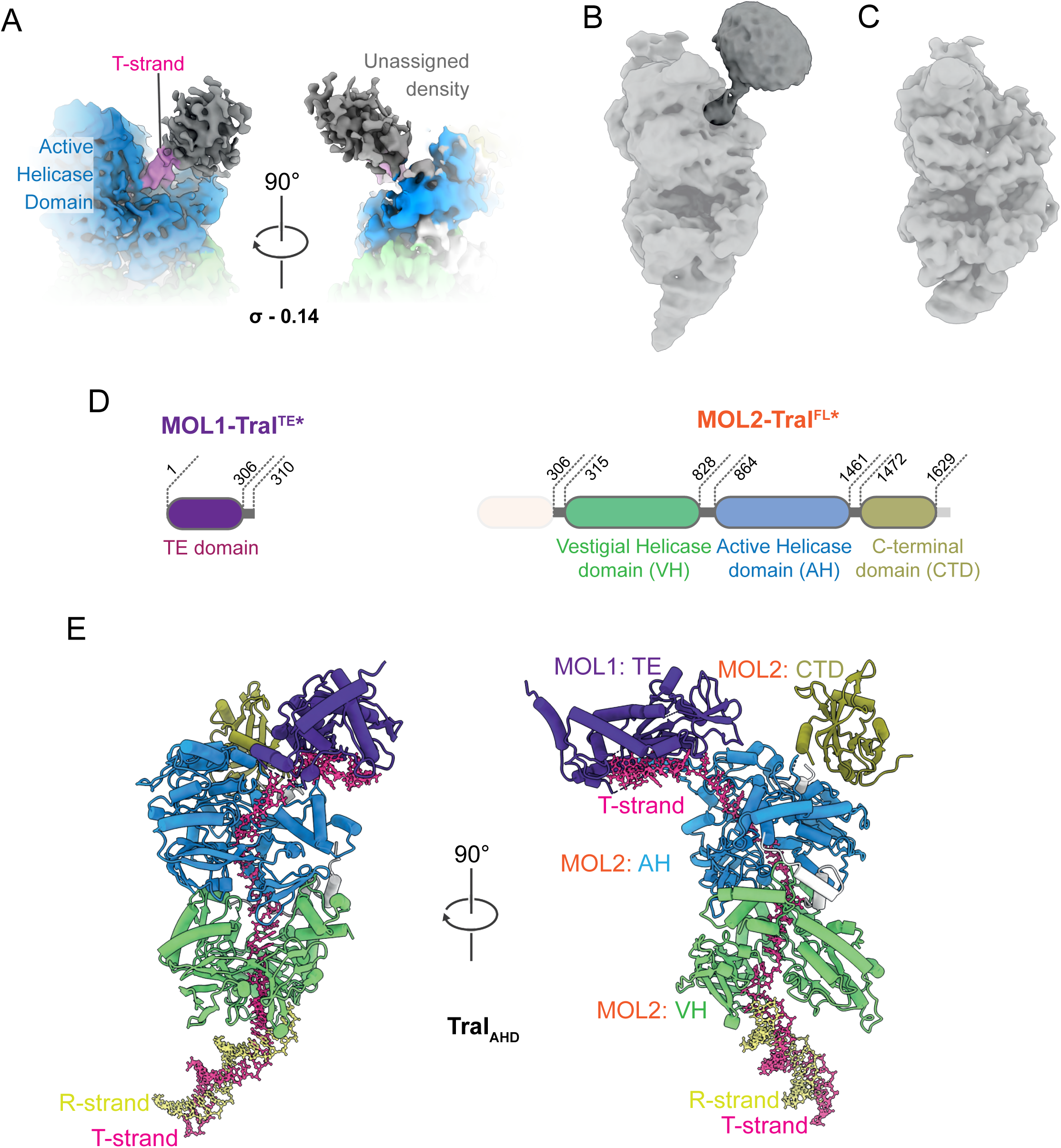
Class-2 represents the functionally asymmetric TraI homodimer (TraIAHD) a. Refined cryo-EM map of class-2 particles, showing additional density at σ = 0.14. b. Cryo-EM map of the tripartite TraI^TE*^:TraI^310-1756^: fDNA65T/26R complex, revealing density for the helicase domains, the forked-DNA substrate, and the TE domain. c. Cryo-EM map of the dipartite TraI^310-1756^:fDNA47T/26R complex, showing density for the helicase domains and the forked-DNA substrate, but lacking detectable TE-domain density. d. Protein constructs used for structural analysis. Left: TraI^TE^ (Mol-1). Right: TraI^FL^ (Mol-2), with only domains that are resolved in the TraIAHD map are highlighted. e. Structure of the TraIAHD complex, in which TraI^TE*^ (Mol-1) and TraI^FL*^ (Mol-2) bind the forked DNA substrate (fDNA65T/26R). Domains visible in the cryo-EM reconstruction are distinctly coloured: TE of Mol-1 (purple); and VH (green), AH (blue), and CTD (olive green) of Mol-2. Two views are shown, rotated by 90°.

To determine the origin of the additional density in state-2, we assembled a new biochemical complex where only one TE domain is present in the sample. This was achieved by employing a TraI construct devoid of the TE domain (TraI^310–1756^) instead of the previously used TraI^FL*^, together with an isolated TE domain (TraI^TE*^) and fDNA65T/26R. This yielded a new tripartite complex (TraI^TE*^:TraI^310–1756^:fDNA65T/26R), containing a TE domain on only one TraI polypeptide, instead of two as before (Figure S1D, S1E and S1F). This new tripartite complex was purified as described and a cryo-EM dataset was collected on this sample representing a positive control for the presence of the TE domain (Figure S1F and S1G). In addition, we employed a TE-lacking negative control, which is a bipartite complex where no TE domains were present, represented by TraI^310–1756^:fDNA47T/26R (Figure S1H and S1I), for which another cryo-EM dataset was obtained (Figure S1J, S1K and S1L).

The new tripartite TraI^TE*^:TraI^310–1756^:fDNA65T/26R complex dataset produced a map closely resembling that of state-2, where additional density attributed to TE was clearly distinguishable (Figure 3B), whereas the map from the bipartite TraI^310–1756^:fDNA47T/26R complex showed no extra density (Figure 3C). The differences between the maps arose from the presence or absence of TE domain in the complex and thus allowed to definitively assign the location of the extra density present in state-2 to the TE domain. Collectively, these data support the notion that the TE domain observed in state-2 does indeed come from a separate molecule and hence the whole assembly must correspond to an asymmetric homodimer of TraI mediated by ssDNA. Therefore, state-2 represents the functionally significant dimer intermediate in the DNA processing pathway. This structure will henceforth be described as TraIAHD, representing the asymmetric homodimer intermediate captured in state-2, within which we attribute TraI^TE*^ as molecule-1 or ‘Mol-1’ and TraI^FL^Y16F/Y17F as molecule-2 or ‘Mol-2’.

Although the TE-domain density was insufficient for an unambiguous rigid-body fit, we could place the TE model tentatively as a rigid body within the low-resolution map. With the help of density exiting the vestigial helicase, ssDNA can be modelled to connect the helicase domains of Mol-2 and TE domain of Mol-1. However, at the current resolution we cannot determine how the ssDNA interacts with the TE domain. Since the active site mutant form of TE (TraI^TE*^ and TraI^FL*^) has been used to reconstitute TraIAHD, we can be sure that no covalent intermediate of TE-ssDNA can be formed in this complex and hence has to represent the TE in the *TE binding site* engaged state, similar to what was observed in previously determined TraI^TE^ crystal structures^15^. Overall, these observations suggest that the complex has two TraI protomers bound to *oriT*; where TraI^TE*^ is engaged to the 5’ side of *nic* at the *TE binding site*, and TraI^FL*^ is bound DNA 3’ to *nic* at the *helicase binding region* (Figure 3D and 3E).

### A single TE domain is involved during initial stages of conjugative DNA processing in the asymmetric TraI dimer

It is intriguing to note that in both state-1 and state-2 reconstructions, only a single TE domain is resolved. In TraIHEL (state-1), the TE density lies in proximity to helicase core and appears to make direct contacts with it, consistent with a TE position tucked adjacent to the helicase motor (Figure 2B). By contrast, in TraIAHD (state-2), the TE density does not make obvious protein-protein contacts with the helicase and sits more at the periphery (Figure 3E). To understand this, we structurally aligned the two maps and compared the orientation of the TE density relative to the TraI helicase modules (Figure 4A). This analysis showed the positioning of TE-domains spatially overlap to a substantial degree when the models are superposed, implying that the two TE domains cannot be accommodated simultaneously without steric clash. This analysis provides a structural rationale explaining why only one TE density is observed in either class. In both TraIHEL and TraIAHD maps, the path taken by the nucleic acid bases supports this interpretation (Figure 2B and 3A). In the TraIHEL map, the ssDNA exiting the helicase tracks toward the TE site, and appears to brush past the TE region, continuing toward the CTD, building upon previous observation^13^ (Figure 4B). In the case of TraIAHD, the ssDNA exiting the helicase (includes the *nic* and the *TE binding* site) is found to be engaging the TE domain of TraI Mol-2, in a manner representing the TE bound configuration (Figure 4C). These observations lead to a working model for the initiation of bacterial conjugation in which two identical TraI protomers adopt non-equivalent, asymmetrical roles. We propose that the TE domain of Mol-1 first engages *oriT*, binding to the *TE binding site*. A second TraI protomer, Mol-2, then loads in its helicase mode onto the *helicase binding region* 3’ to *nic*. At this stage, the TE of Mol-2 remains disengaged from *oriT*, because the TE site is already occupied by Mol-1, and the two TE positions are mutually exclusive at this stage (Figure 4D).

**Figure 4.**
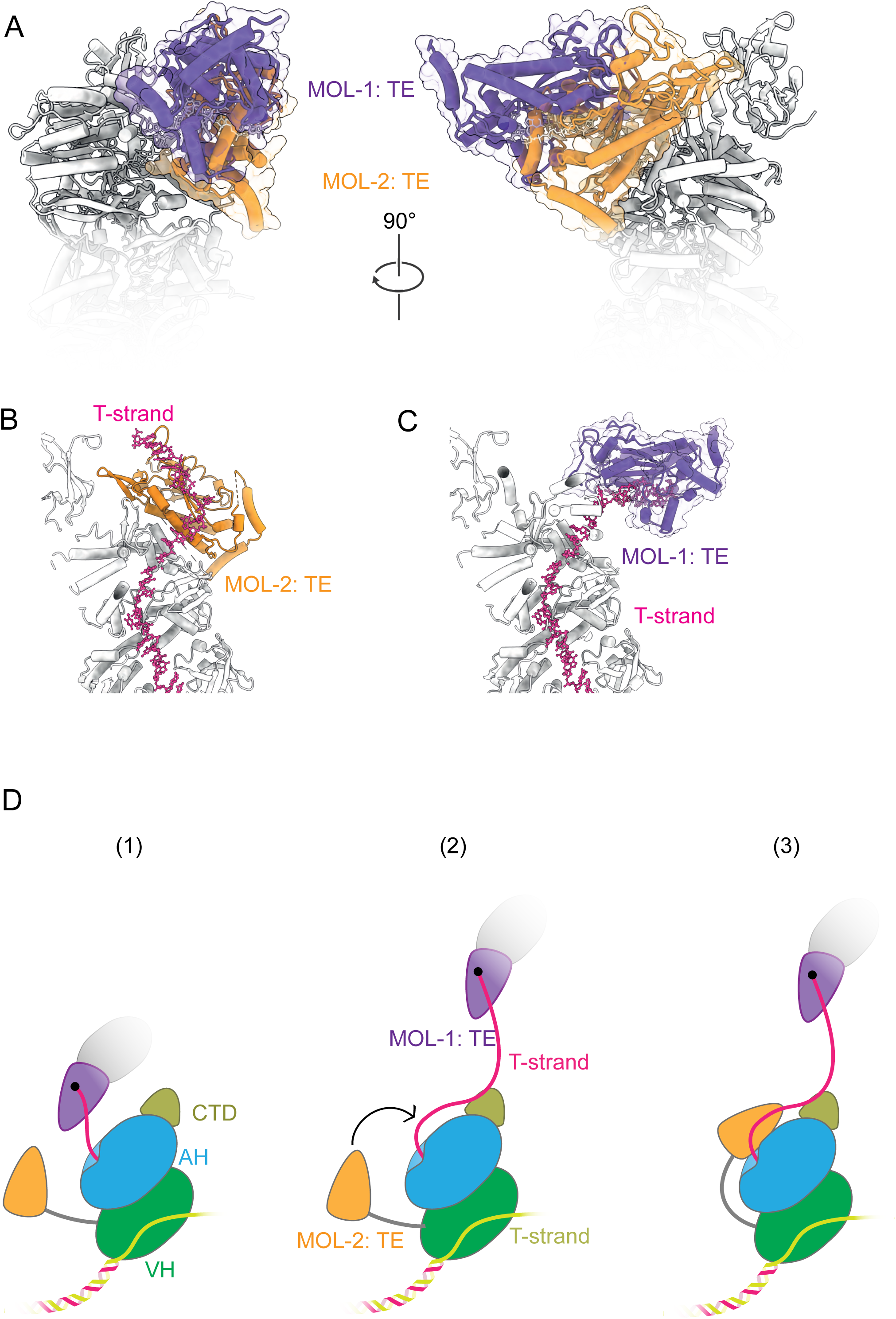
Structural comparison of TraIHEL and TraIAHD. a. Structural alignment of the TraIHEL and TraIAHD complexes. The TE domains are highlighted: the TE domain from TraIHEL is shown in orange, and the TE domain from TraIAHD in purple. All other regions of TraI are shown in light grey. b. The TE domain of TraIHEL (orange) engaging the T-strand ssDNA (pink) as it exits the helicase core. c. The TE domain of TraI(TE) Mol1 (purple) in the TraIAHD complex engaging the T-strand ssDNA (pink) exiting the helicase core. d. Schematic model illustrating TE-domain engagement with the T-strand ssDNA at different stages of DNA processing. (i) The TraIAHD state where the TE domain of Mol-1 is engaged with the T-strand, while the TE domain of Mol-2 remains disengaged. (ii) After initial ssDNA processing, the TE domain of Mol-1 may be released, potentially piloting T-strand transfer. (iii) The TE domain of Mol-2 to engage the ssDNA and transition the complex into the TraIHEL state.

### TraI helicase prefers to load onto an unnicked supercoiled DNA template

Since the cryo-EM structures described here were determined on a synthetic forked DNA substrate emulating a pre-opened duplex (Figure 1C), while the native substrate is fully closed supercoiled DNA (scDNA), we next asked how the asymmetric homodimer of TraI loads onto scDNA substrate. The TE domain has been shown to be nicking-competent on scDNA substrate when present in excess and by-passes the need for the relaxosome components to facilitate TE loading onto its cognate sequence within *oriT* (Figure S7A). Prior work has shown that the TraI helicase in its ssDNA-engaged closed state is protected from trypsin proteolysis, whereas TraI in an ssDNA-disengaged or TE-bound, open state is readily cleaved^13^. We therefore used the trypsin proteolysis assay to report on helicase loading on scDNA. The condition involving the TraI^310–1756^ construct (which lacks a TE domain) alone and no TraI^TE^ available *in-trans*, showed minimal protection, indicating poor intrinsic loading on scDNA by TraI’s helicase domains themselves, independent of TE. When TraI^310–1756^ was supplemented with TraI^TE*^, we observed increased protection from trypsin proteolysis, evident as a high–molecular-weight band corresponding to protected TraI^310–1756^, consistent with helicase loading. In contrast, replacing TraI^TE*^ with wild-type TraI^TE^ abolished this protected band, indicating that helicase loading does not happen on nicked scDNA (Figure 5A). We repeated the assay substituting TraI^310–1756^ with TraI^FL*^ and obtained similar results across conditions apart from the expected difference in the TraI^FL*^ alone condition. In the TraI^FL*^ alone condition, protection against trypsin proteolysis can be observed, suggesting TraI helicase loading onto the scDNA. The TraI^FL*^ construct includes the TE domain necessary for initial binding, enabling subsequent helicase loading of TraI helicase domains that are present in stoichiometric excess (Figure 5B). Taken together, these experiments indicate that the TE domain must be present to enable helicase loading. We can deduce that TE engages its cognate site first, creating the conditions favourable for helicase engagement. Crucially this assay highlights that when the DNA substrate is nicked, TraI’s helicase domains do not load, showing that helicase loading requires an unnicked scDNA substrate.

**Figure 5.**
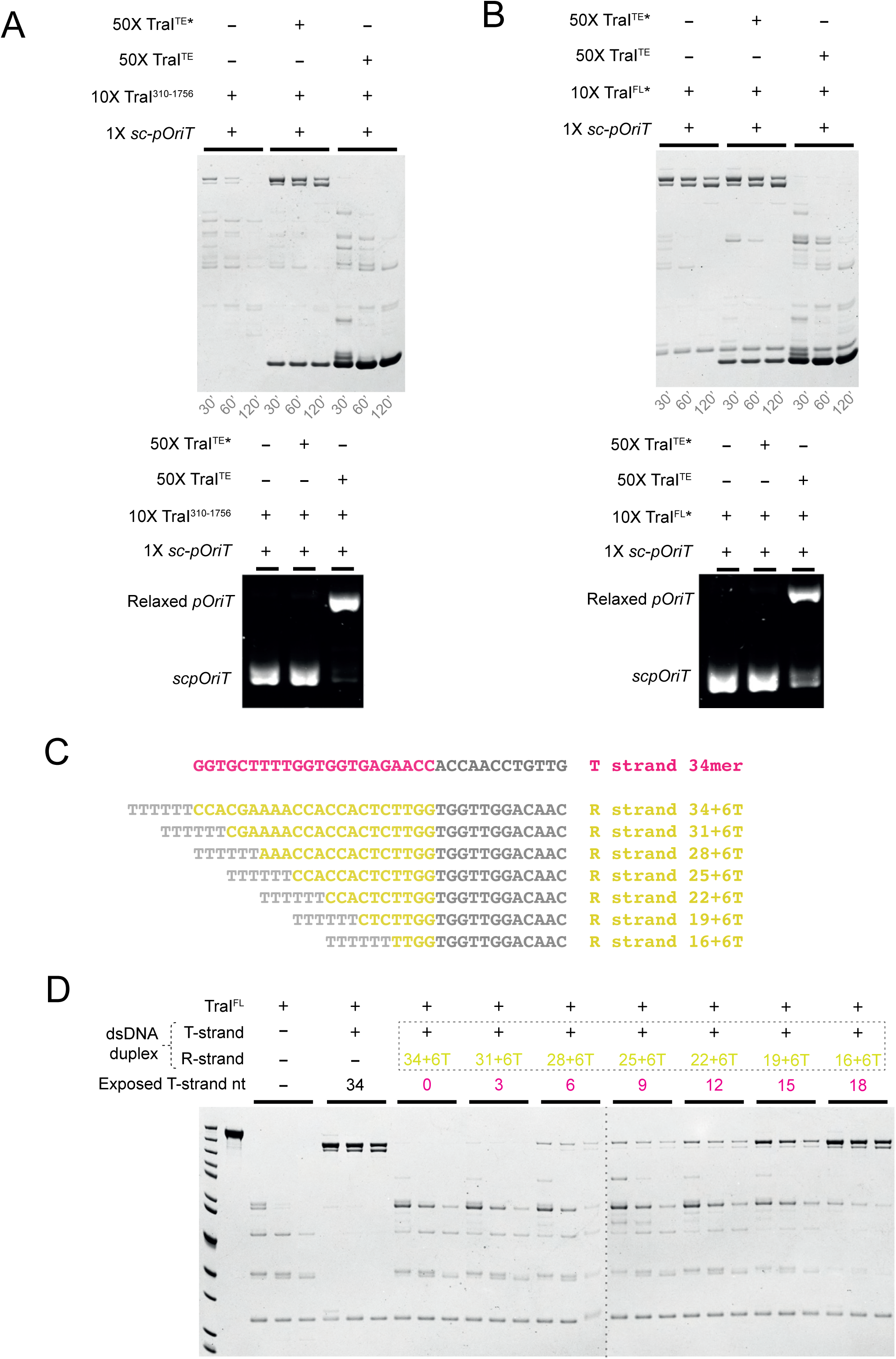
Conditions governing the loading of TraI as a helicase. a. Top - Trypsin proteolysis assay probing the conformation of TraI^310-1756^ bound to supercoiled plasmid DNA (*scporiT*) under three conditions: in the absence of TE or TE*, in the presence of TE*, or in the presence of wild-type TE. Samples were digested with trypsin for 30, 60, or 120 minutes and analysed by SDS–PAGE as described in Methods. Bottom – Agarose gel of the same sample after proteinase K treatment, showing the unnicked *scporiT* and relaxase plasmid with *oriT*. b. Top – Trypsin proteolysis assay probing the conformation of TraI^FL^ bound to *scporiT* under the same three conditions described in panel (a). Samples were digested for 30, 60, or 120 minutes and analysed by SDS–PAGE. Bottom – Agarose gel of the same sample after proteinase K treatment, showing the unnicked *sc-poriT* and relaxed plasmid with *oriT*. c. Sequence of the T-strand (34-mer) oligonucleotide used in panel (d), along with the series of complementary R-strand oligonucleotides designed to generate DNA fork substrates exposing progressively longer stretches of T-strand ssDNA. d. Trypsin proteolysis assay examining the conformation of TraI^FL^ bound to DNA fork substrates that expose increasing lengths of T-strand ssDNA (3 to 18 nucleotides). Each TraI– DNA complex was digested with trypsin for 30, 60, or 120 minutes and analysed by SDS– PAGE.

If TraI preferentially loads as a helicase on an unnicked template, a local ssDNA bubble must therefore form within the native supercoiled DNA substrate to permit helicase engagement. Hence, we asked what the minimum length of exposed ssDNA (the T-strand) is required to support TraI helicase loading. To probe this, we designed a forked DNA series derived from *oriT* to titrate the extent of ssDNA exposure on the T-strand. The T-strand was a 34-mer comprising the established helicase-binding site (22mer) plus 12 additional nucleotides, to enable strand annealing. The complementary R-strand was systematically truncated from its 3′ end in 3-nt steps, incrementally exposing between 0 and 18 bases of the T-strand. A short poly-T tail was appended to the 3′ end of the R strand to create a fork (Figure 5C). We used limited trypsin proteolysis as a readout of the helicase “closed” state (protection from proteolysis), with the 34-mer T-strand alone containing the fully exposed helicase binding site and therefore serving as a positive control for helicase loading. With the DNA fork substrates, both the 0 and 3 base (T-strand) exposed conditions showed no protection, indicating the absence of the closed helicase state. By contrast, protection became detectable when as little as 6 bases of the T-strand were exposed and increased progressively with the extent of ssDNA opening, approaching the protection seen with the 34-mer ssDNA control (Figure 5D). These data indicate that while TraI prefers longer stretches of exposed T-strand ssDNA, it can still adopt the closed, helicase-loaded state with as few as 6 T-strand nucleotides exposed. Thus, formation of an expansive ssDNA bubble is not required for helicase loading as previously suggested^16^. Indeed, a larger bubble would likely be unstable and energetically unfavourable under physiological conditions.

### TraI unwinds dsDNA in an ATP independent manner to load as a functional helicase

TraI has been reported to bind 18nt of ssDNA to adopt its closed, helicase mode conformation as suggested by structural work reported here and earlier^13^. However, our biochemical analysis in Figure 5D show that TraI can attain this closed state with as little as 6 exposed nucleotide bases on the T-strand. This prompted us to ask how TraI loads on a mixed ds/ss duplex template. We first used a DNA fork substrate comprising a 22 nt T-strand that includes the sequence of the *helicase binding region*, with 11 nt exposed and 11 nt base-paired to an R-strand; a short poly-T extension at the R-strand 3′ end created the fork (fDNA22T/11R). The T-strand was FITC-labelled at its 3’ end, while the R-strand carried a Cy5 label at the 3’ end (Figure 6A). TraI^FL^ was titrated at 0.2, 0.5, 1, 2, and 4 × molar excesses over the DNA substrate and monitored by gel-shift. We observed an increase in the FITC signal in shifted species concomitant with a reduction of the duplex band’s intensity. Additionally, the appearance of a free Cy5-R-strand is consistent with TraI selectively engaging the T-strand and unwinding the 11-bp duplex segment (Figure 6B). To test the involvement of ATP, we repeated the assay with the previously described helicase activity deficient mutant TraI^K998M^ (Walker-A, ATP-binding–deficient) with and without ATP conditions^15,17^. The results were the same as with the wildtype, indicating that TraI recognises the 11 exposed T-strand nucleotides and separates a further ~11 bp as it folds into its closed helicase conformation in an ATP-independent manner (Figure 6C and 6D). We next asked how far this ATP-independent strand separation extends. To probe this, a longer DNA fork substrate containing a 5’-FAM labelled 34 base T-strand was implemented, in which 11 bases are exposed and 23 nucleotides are base paired to the 3’ Cy5-labelled R-strand (fDNA34T/23R) (Figure 6E). A titration of TraI at 0.2, 0.5, 1, 2, and 4 × molar excesses over the fDNA substrate shows that with an increasing concentration of TraI^FL^ and TraI^K998M^, the fDNA34T/23R band, disappeared but no free R-strand appeared. Instead, both FITC- and Cy5-signals co-migrated together with TraI^FL^ and TraI^K998M^ in a slower-moving complex (Figure 6F and 6G). This indicates that beyond the initial local strand separation seen with shorter annealing sites, TraI remained bound to the T/R duplex DNA segment but was rendered incapable of unwinding it in the absence of ATP (Figure 6H). When ATP was included as a control, TraI^FL^ promoted further unwinding (Figure S6A and S6C), whereas TraI^K998M^ did not (Figure S6B and S6D). Together, these data suggest that TraI can load as a helicase by recognising a short, exposed ssDNA patch of the T-strand and locally separate additional base pairs to achieve the closed helicase state without ATP. An expansive ssDNA bubble is neither necessary nor favoured at the loading step; once loaded as a helicase, TraI unwinds DNA in an ATP-dependent manner, consistent with previous observations^18^.

**Figure 6.**
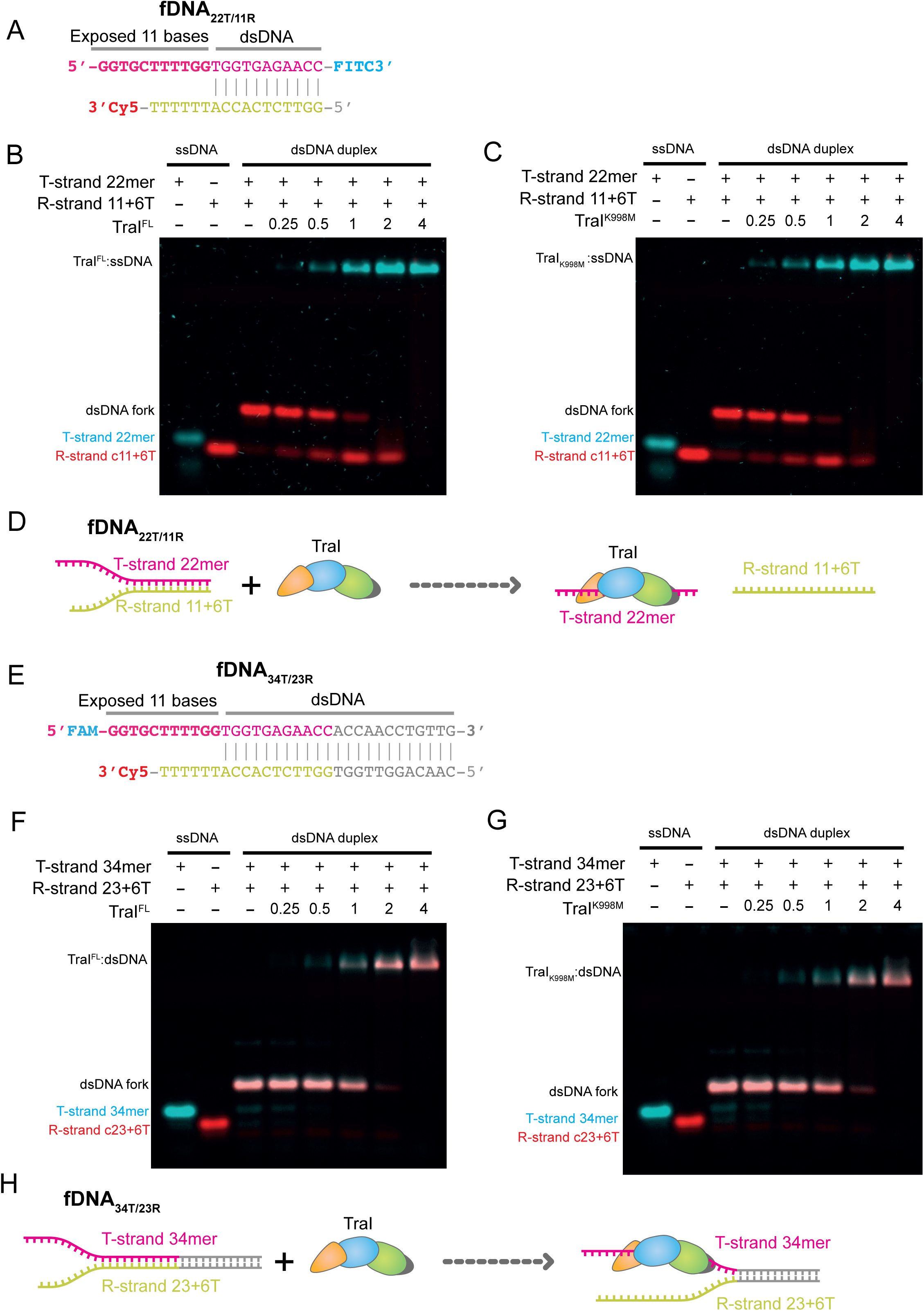
ATP-independent loading of TraI as a helicase on minimally exposed forked DNA. a. Sequence of the fDNA22T/11R template used to assess TraI helicase loading. In the duplex region, the T-strand is labelled at its 3′ end with FITC and the R-strand with Cy5. b. Electrophoretic mobility shift assay (EMSA) examining binding of TraI^FL^ to the 22-mer T-strand, 16-mer R-strand, or the duplex form of fDNA22T/11R. TraI was titrated at 0.25×, 0.5×, 1×, 2×, and 4× molar equivalents relative to the DNA substrate. Experimental details are provided in Methods. c. EMSA as in panel (b), using the ATPase-deficient TraI^K998M^ variant. d. Schematic summary illustrating TraI loading on the fDNA22T/11R substrate. e. Sequence of the fDNA34T/23R template used for helicase-loading assays. The T-strand is labelled at the 5′ end with FAM, and the R-strand at the 3′ end with Cy5. f. EMSA examining binding of TraI^FL^ to the 34-mer T-strand, 29-mer R-strand, or the duplex fDNA34T/23R substrate. TraI was titrated at 0.25×, 0.5×, 1×, 2×, and 4× molar equivalents. g. EMSA as in panel (f), performed with TraI^K998M^. h. Schematic summary of TraI loading on the fDNA34T/23R substrate.

### The relaxosome protein IHF inhibits the TE mediated nicking activity of TraI Mol-1

Our biochemical work reported here demonstrated that TraI preferentially loads as a helicase onto an unnicked DNA substrate (Figure 5A and 5B). Further, these results suggest the order of assembly of the asymmetric TraI homodimer. The TE domain of TraI Mol-1 binds *oriT* first, thereby enabling the subsequent loading of TraI Mol-2 as a helicase. This raises the key mechanistic questions of how Mol-1 TE-domain loading does not immediately trigger nicking, and how the plasmid is maintained in an unnicked state long enough to permit productive helicase loading.

Investigating this further, we noticed that IHF has its binding site closest to the *TE binding site* in *oriT*. We questioned if IHF could play a role in inhibiting the nicking activity of Mol-1 TE due to its proximity to the TE domain within *oriT*. To test whether IHF inhibits TE-mediated nicking, we first assayed nicking of *sc-poriT* at a fixed DNA concentration in the presence or absence of a 10-fold molar excess of IHF, varying TraI^TE^ over 1 ×, 2×, 10 ×, 50 × and 100 ×. At 1 × and 2 × molar excesses of TraI^TE^, no appreciable nicking was detected with or without IHF. At 10 × or higher molar excesses of TraI^TE^, robust nicking was observed only in the absence of IHF, while no nicking occurred when IHF was present (Figure S7A). We then examined full-length TraI (TraI^FL^) under similar conditions and as reported previously^19^, IHF no longer suppressed nicking by TraI^FL^, an effect that was particularly evident at higher molar excesses of TraI^FL^ (Figure S7B). Given that TraI can only load as a helicase onto an unnicked DNA template, with this loading requiring the prior loading of TE-domain (Mol-1) onto its cognate binding site (*TE binding site*), we delineated the two activities by reconstituting the TE and helicase domains separately via TraI^TE^ and TraI^310-1756^ constructs. TraI^TE^ was able to nick *sc-poriT* and this nicking was indeed inhibited in the presence of IHF. Remarkably this IHF mediated inhibition was abolished in the presence of TraI^310–1756^ (Figure S7C). These findings suggest that IHF inhibits TE mediated nicking, thus enabling TraI loading as a helicase, which can only load on unnicked template. TraI loading as a helicase eventually releases the IHF mediated nicking inhibition of TE, providing a regulatory loop for the initiation of conjugative DNA processing mechanism.

## Discussion

Conjugative plasmid transfer is the major route for horizontal gene dissemination, yet the mechanistic details of how the relaxase (TraI) coordinates origin processing have remained poorly defined. While it is established that TraI carries both trans-esterase and helicase functions, the temporal order of their engagement, the nature of the dimeric complex involved, and the regulation of nicking at *oriT* were unclear^12,13^. Moreover, the structural basis for how two identical TraI protomers partition functionally distinct roles has not been resolved. In this study, we address these longstanding questions by identifying and characterizing a key intermediate in the conjugation pathway, a functionally asymmetric TraI homodimer assembled on *oriT*. Our work provides the first structural insight into this key intermediate in which two TraI molecules assemble a functionally asymmetric homodimer. Notably, the two TraI protomers do not engage in direct protein-protein interactions; rather, their asymmetry arises entirely from distinct engagements with DNA. This DNA-mediated architecture explains how identical subunits adopt sequential, non-equivalent roles: one molecule (Mol-1) executes the trans-esterase reaction, while a second (Mol-2) loads as a helicase to initiate strand separation.

Although the structure of TraI in its closed helicase state has been described previously, the mechanism of duplex DNA unwinding remained unresolved^13^. Earlier work employed ssDNA substrates and could not address how TraI engages and separates DNA duplex. In RecD2, an archetypal SF1B helicase and structural homologue of TraI, strand separation is mediated by a β-hairpin loop that acts as a strand-separation wedge^20^. While our structural data reveal a similar β-hairpin loop near the duplex junction, it does not appear to function in the same way^13^. Mutagenesis of key residues in this loop failed to impair conjugative transfer, suggesting that TraI employs a distinct mechanism for duplex unwinding.

Traditionally, DNA processing during conjugation was assumed to mirror rolling-circle replication. In that canonical model, an initiator protein (e.g., Rep) first binds and nicks the origin of replication, after which a helicase (e.g. PcrA) is recruited^21,22^. However, our data overturn this assumption by demonstrating that TraI loads as a helicase on an unnicked *oriT* template. This implies that helicase recruitment must occur on a transiently destabilised duplex, potentially involving the formation of a small DNA bubble. Earlier structural studies show that TraI binds approximately 18 nucleotides of ssDNA in its helicase mode, raising the question of how such a region becomes accessible in the absence of prior nicking^13^. A comparable step in bacterial replication initiation relies on a more expansive mechanism, where multiple copies of the master replication initiator DnaA cooperatively opens duplex DNA to expose ssDNA and thereby enables subsequent helicase loading^23^. In contrast, TraI employs only a single TE domain to recruit the helicase-competent TraI protomer. This led us to systematically define the minimal ssDNA region required for TraI helicase loading. Remarkably, we found that as few as six nucleotides are sufficient, although longer exposures likely improve efficiency. Strikingly, once bound to a forked substrate with minimal ssDNA (11 – T-strand), TraI itself promotes further DNA duplex melting without ATP hydrolysis, to the complete extent of helicase loading. Though this phenomenon of ATP independent strand separation has been observed in helicases such as NS3 and Pif1, the driving force or energy source to mediate duplex unwinding is unclear^24,25^.

While our findings suggest that TraI Mol-1 is loaded first via its TE domain, with helicase engagement by TraI Mol-2 occurring subsequently, it remains unclear how an ostensibly nicking-competent TE domain can be poised at *oriT* without cleaving it. In F-family *oriTs*, two well-defined IHF binding sites are typically present, with *IHF-A* located immediately upstream of the *TE binding site*^9^. IHF is a heterodimeric, chromosome-encoded protein that introduces sharp (~160ᵒ) DNA bends^26^ and plays major roles in the host cell including transcriptional regulation and genomic structural remodelling^27^. Its constitutive presence in the bacterial cytoplasm presents it as a major regulator in the host^28,29^. Previous work has shown that IHF enhances the nicking activity of TraI in the F and R1 plasmids while inhibiting nicking in the R388 systems^19,30^. Our findings indicate that IHF inhibits nicking when the TE domain is present alone (TraI^TE^), but not when full-length TraI is present, a result consistent with prior observations in F and R1 plasmid systems^19,31^. By systematically probing the effects of separate TE and helicase domains of TraI, it became evident that the helicase domain relieves the IHF-mediated inhibition of TE-catalysed nicking (Figure S7).

To rationalize these observations, we considered the recently published F-plasmid relaxosome structure^11^, in which IHF engages the vestigial helicase domain of TraI (TraI^VH^) but makes no direct contact with the TE domain. In this complex, the TE domain is positioned on its cognate *oriT* site (*TE binding site*) and corresponds to TraI Mol-1 in our model. Our biochemical data show that IHF inhibits the nicking activity of the TE domain when it is functionally isolated (TraI^TE^), but this inhibition is relieved when the helicase domains, particularly the VH domain is available, either *in cis* (TraI^FL^; Figure S7B) or *in trans* (TraI^310–1756^; Figure S7C). Together, we use these findings to argue that the IHF–TraI^VH^ interaction observed in the F-plasmid relaxosome is unlikely to inhibit TE-mediated *oriT* nicking; instead, it likely stabilizes a nicking-competent configuration of the TE domain. We therefore propose that an earlier intermediate must exist, in which the TE domain of TraI Mol-1 is retained in a non-nicking-competent configuration prior to productive engagement of its cognate binding site. Previous work has suggested that TE reaches the *TE binding site* via a pre-exposed T-strand bubble 3′ to the *nic* site that allows the TE domain to slide into the binding site^11^. It is possible that IHF prevents full access of the TE domain to its cognate *TE binding site*, either directly or indirectly by reshaping *oriT* around *nic*, thereby gating TE engagement and delaying nicking until helicase loading by TraI Mol-2 occurs.

Based on our structural and biochemical data, we propose the following model for conjugation initiation. The relaxosome accessory proteins are bound to *oriT*, then TraI Mol-1 gains partial access via its TE domain to its cognate binding site. IHF keeps the TE domain of TraI Mol-1 in a non-nicking configuration, however this event destabilises the duplex and exposes a small stretch of nucleotides 3′ of *nic*, thereby creating the ssDNA patch sufficient to load TraI Mol-2 as a helicase. IHF bound adjacent to the *TE binding site* maintains the *oriT* in an unnicked state, extending the window for helicase engagement. Mol-2 then loads onto this transient ssDNA patch as a helicase. In this scheme, IHF imposes a regulatory brake on the TE activity of Mol-1, so that helicase loading is coordinated to precede nicking. Once Mol-2 is loaded as a helicase it relieves IHF-mediated nicking repression of Mol-1, coupling helicase loading to initiation of strand transfer. With IHF displaced, the TE domain of Mol-1 then nicks the T-strand, forming the covalent phosphotyrosyl intermediate that marks the onset of transfer. As unwinding proceeds, the nucleoprotein assembly is remodelled, favouring Mol-1 disengagement. The T-strand exiting the Mol-2 is then captured by its own TE domain, consistent with the observed ssDNA trajectory and single TE occupancy in both structural classes (Figure 2B and 3E). This dynamic, DNA-mediated division of labour - one TE-active and a second helicase active - provides a unified mechanistic basis for the functional asymmetry seen in our maps and reconciles it with the biochemical reconstitution experiments (Figure 7).

**Figure 7:**
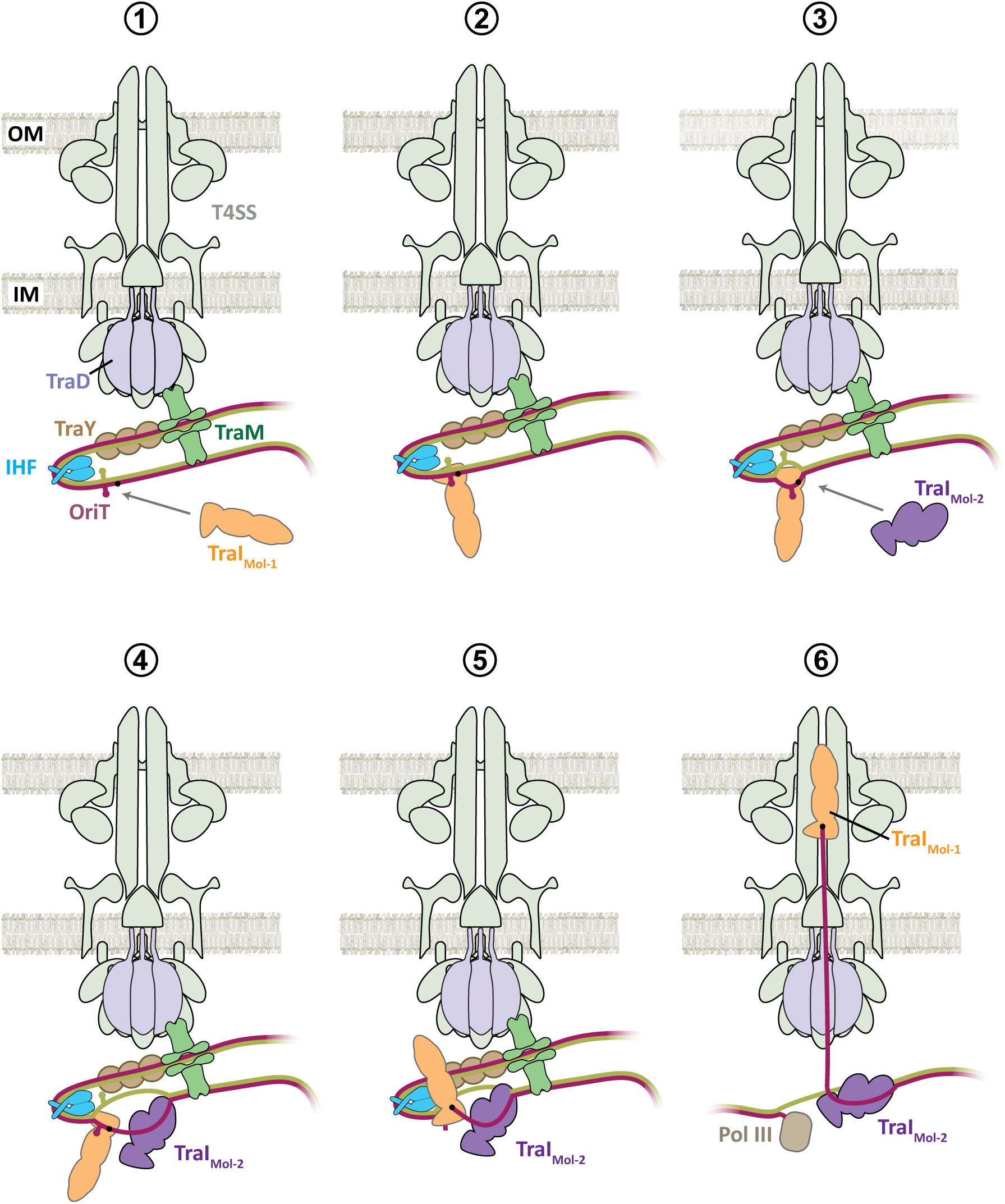
Schematic model of DNA processing steps mediated by two molecules TraI prior to initiation of conjugative DNA transfer. This model incorporates current findings together with past findings and provide a mechanistic overview for the initiation of conjugation and ultimately for the spread of antibiotics resistance genes in bacteria. Step-1: Accessory proteins are pre-bound at the relaxosome, poised to receive TraI Mol-1. Step-2: The TE domain of Mol-1 gains partial access to its *oriT* binding site, while *oriT* remains unnicked due to IHF-mediated inhibition. Step-3: The TE domain of Mol-1 destabilises the local duplex and exposes a small number of T-strand bases, enabling TraI Mol-2 to load as a helicase. Step-4: Using these exposed T-strand bases, TraI Mol-2 extends unwinding to generate the longer ssDNA region required for stable helicase loading, 3′ of the *nic*. Step-5: Helicase loading by Mol-2 triggers a conformational change in Mol-1 that relieves IHF-mediated nicking inhibition, allowing the TE domain of Mol-1 to nick *oriT* and form the covalent nucleoprotein intermediate. Step-6: This initiates conjugative DNA transfer: Mol-1 remains associated with and pilots the transferred T-strand, while Mol-2 continues unwinding plasmid DNA to supply ssDNA for transfer.

By redefining the sequence of initiation steps and uncovering a previously unappreciated regulatory checkpoint imposed by IHF, our findings fundamentally revise the model of conjugative transfer initiation. This mechanism, in which helicase loading precedes nicking and remodels the DNA–protein complex to relieve repression, offers a new framework for understanding relaxosome dynamics. The requirement for functional asymmetry, minimal ssDNA exposure, and ATP-independent DNA melting highlights a tightly controlled initiation switch that could be selectively targeted to block horizontal gene transfer and combat the spread of antibiotic resistance.

## Methods

### Cloning, expression and purification of TraIFL, TraI^TE^, TraI^310-1756^ and their mutants

Mutant and deletion constructs were generated from the TraI^FL^ construct previously described^13^ and Platinum SuperFi II DNA Polymerase (Invitrogen). DNA fragments were assembled into the pCDF-1b expression vector (Novagen), under control of a lac promoter, using In-Fusion® Snap Assembly (Takara Bio). TraI^FL^ and TraI^310-1756^ contained an N-terminal His6-tag followed by HRV-3C protease cleavage site, which was not removed during purification. TraI^TE^ contained a C-terminal Strep-tag II.

For *in vivo conjugation assays*, the *traI* mutants were generated via In-Fusion cloning to introduce specific point mutations. Primers were designed to harbour the desired amino acid substitutions flanked by 15-bp overlapping sequences homologous to the pCDF-TraI vector backbone. The target plasmid was linearized by PCR, and the mutated fragments were directionally assembled using the In-Fusion enzyme suite. Following transformation into *E. coli* Top10 cells, successful site-directed mutagenesis and the integrity of the *traI* gene were confirmed by DNA sequencing.

The R1ΔtraI plasmid was constructed using classical one-step gene inactivation via FLP–FRT recombination. Briefly, *E. coli* Top10 cells harbouring the R1 plasmid carrying a kanamycin resistance cassette were transformed with pWRG99 (Table S4), which expresses the λ Red recombinase system, and plated on LB agar supplemented with 100 µg mL^−1^ ampicillin at 30 °C for two days. An overnight culture was diluted 1:20 in fresh LB and grown at 30 °C for 1 h, after which λ Red expression was induced by addition of L-arabinose to a final concentration of 0.1%. When cultures reached an OD_600_ of 0.7, cells were harvested and made electrocompetent. Primers I and II (Table S5) and plasmid pKD3 (Table S4) were used to amplify an FRT-flanked chloramphenicol resistance cassette with 50-bp homology arms corresponding to the regions immediately upstream and downstream of the *traI* open reading frame. The PCR product was treated with DpnI, purified, and electroporated into the prepared cells, which were subsequently selected on LB agar containing 50 µg mL^−1^ kanamycin and 25 µg mL^−1^ chloramphenicol and incubated overnight at 37 °C. Successful recombinants were screened by colony PCR using primers III and IV (Table S5). Correct clones were then transformed with pCP20 (Table S4), which encodes the Flp recombinase, and plated on LB agar supplemented with 100 µg mL^−1^ ampicillin at 30 °C for two days. Transformants were grown overnight in liquid culture at 30 °C, subcultured into fresh LB, and incubated at 42 °C for 5 h to cure the temperature-sensitive plasmid and excise the resistance cassette. Serial dilutions were plated on LB agar containing 50 µg mL^−1^ kanamycin and incubated for 24 h at 37 °C. Colonies were screened for loss of chloramphenicol resistance by replica plating, and the resulting clones were verified by sequencing to confirm deletion of the *traI* locus. The lists of plasmids used in the conjugation assay and in the structural/biochemical experiments are provided in Tables S1 and S2, respectively.

All TraI constructs (TraI^FL^, TraI^310-1756^, TraI^TE^, TraI^K998M^, TraI^FL^ and TraI^TE^) were expressed and purified with a similar workflow implemented for TraI^FL^ that was described previously^13^. Briefly, recombinant plasmids containing tagged TraI constructs were transformed into *E. coli* BL21 star (DE3) cells (Thermo Fisher Scientific). Cells were grown in LB medium supplemented with 100 µg/ml spectinomycin at 37 °C to an OD600 of 0.6–0.8. Expression was induced with 1 mM IPTG, and cultures were incubated for 16 h at 17 °C. Cells were harvested by centrifugation and resuspended in lysis buffer (50 mM Tris-HCl pH 7.5, 100 mM NaCl, 5% (v/v) glycerol, protease inhibitor cocktail (Roche)). Lysates were clarified by centrifugation at 38,000 × g for 60 min, and the soluble supernatant subjected to FPLC purification.

TraI^FL^ and TraI^310-1756^ constructs were first purified by Ni^2+^ affinity chromatography using a 5 ml HisTrap HP column (Cytiva), eluting over an imidazole gradient (25–500 mM). TraI^TE^ constructs were purified by Strep-Tactin affinity chromatography using a 5 ml Streptrap HP column (Cytiva), eluting with lysis buffer supplemented with 2.5 mM D-desthiobiotin (Sigma Aldrich). Fractions containing TraI were pooled, buffer exchanged into 50 mM Tris-HCl pH 7.5, 50 mM NaCl, 5% (v/v) glycerol, and further purified by anion exchange chromatography using a 5 ml HiTrap Q column (Cytiva) with a 50 mM–1 M NaCl gradient. The final purification step was size exclusion chromatography in 50 mM Tris-HCl pH 7.5, 100 mM NaCl, 5% (v/v) glycerol, using a Superdex 75 10/300 column (Cytiva) for TraI^TE^ and a Superdex 200 Increase 10/300 column (Cytiva) for TraI^FL^ and TraI^310-1756^. Purified proteins were concentrated, analysed by SDS-PAGE, flash frozen in liquid N2 and stored at −80 °C.

### Assembly and purification of the TraI^TE^:TraI^FL^:fDNA; TraI^TE^:TraI^310-1756^:fDNA; TraI^310-1756^:fDNAs complex

DNA oligonucleotides for the T-strand (65 nt; containing the TraI helicase and transesterase binding sites) and the complementary R-strand (34 nt) (Eurofins Genomics) were resuspended to 100 µM in oligonucleotide buffer (20 mM Tris-HCl pH 7.2, 100 mM NaCl, 0.1 mM EDTA). The T-strand was mixed with a 1.2-fold molar excess of R-strand, heated to 95 °C for 20 min, and slowly cooled to room temperature to anneal and generate a forked DNA substrate (fDNA65T/26R). Samples were clarified by centrifugation (11,000 × g, 10 min) and applied to a Superdex 200 Increase 10/300 GL column (Cytiva) pre-equilibrated in complex buffer (50 mM Tris-HCl pH 7.5, 100 mM NaCl, 10 mM MgCl₂). Fractions corresponding to the annealed fork were pooled and quantified on a NanoDrop spectrophotometer (Thermo Fisher Scientific).

The tripartite TraI^TE*^:TraI^FL*^:fDNA65T/26R and TraI^TE*^:TraI^310-1756^:fDNA65T/26R complexes, were assembled at a 15:1:10 molar ratio (TraI^TE*^:fDNA65T/26R:TraI*), using either TraI^FL*^ or TraI^310–1756^ as TraI* for the respective complex. For the bipartite TraI^310–1756^:fDNA47T/26R complex, TraI^310–1756^ and fDNA47T/26R were mixed at a 1:1.2 molar ratio. All reactions were incubated for 30 min at 25 °C and purified on a Superdex 200 Increase 3.2/300 GL column (Cytiva) equilibrated in complex buffer. Peak fractions were analysed by SDS–PAGE.

### Cryo-EM sample preparation and data collection

For all complexes subjected to cryo-EM analysis in this study, Quantifoil™ Au R2/2 200 holey carbon mesh grids were glow-discharged in air at 0.2 bar using a Harrick plasma cleaner (30 s at low intensity, 60 s at medium intensity). The purified complexes (4 µl) were applied on the grids, then blotted for 2.5–2.7 s at 4 °C and 100% humidity, and plunge-frozen in liquid ethane at −196 °C using a Leica EM GP2 automatic plunge freezer. Grids were screened for ice thickness and particle dispersion using a JEOL JEM-2100Plus microscope operating at 200 kV with a LaB6 gun and OneView camera (Gatan). Grids similar to the condition screened were used for data collection on a Titan Krios G3i (Thermo Fisher Scientific) operating at 300 kV with an XFEG G2 gun and Bioquantum K3 direct electron detector (Gatan) at LonCEM, London. Micrograph movies were acquired over a defocus range of −0.5 to − 3.0 µm. For the TraI^TE*^:TraI^FL*^:fDNA65T/26R and TraI^TE*^:TraI^310-1756^:fDNA65T/26R complexes, movies were recorded in electron-counting mode at a nominal magnification corresponding to a calibrated pixel size of 0.84 Å, with a total electron dose of 63 e^−^/Å^2^ and 67 e^−^/Å^2^ respectively, fractionated over 50 frames for each complex. For the TraI^310–1756^:fDNA47T/26R complex, a calibrated pixel size of 0.84 Å, with a total electron dose of 60 e^−^/Å^2^ fractionated over 50 frames was used. Automated acquisition was performed using EPU v3.4 (Thermo Fisher Scientific). Two datasets of 5,964 and 28,157 movies were collected for the TraI^TE*^:TraI^FL*^:fDNA65T/26R complex; 14,594 movies for the TraI^TE*^:TraI^310-1756^:fDNA65T/26R complex, and 13,756 movies for the TraI^310–1756^:fDNA47T/26R complex.

### Cryo-EM image processing and reconstruction

For all datasets movie frames were aligned and corrected for beam-induced motion on-the-fly using MotionCor2^32^. 34,121 corrected and aligned micrographs were imported and all downstream processing was carried out using CryoSPARC v3.3.2 unless specifically stated^33^. For the TraI^TE*^:TraI^FL*^:fDNA65T/26R complex dataset, CTF parameters were estimated using Patch CTF and 25,058 micrographs were selected for further processing based on ice thickness, CTF-fit and astigmatism. An initial set of particles was obtained from a subset of 522 micrographs by blob picking using the circular and elliptical blob function, covering a particle diameter between 90 and 150 Å. Further sorting of particles through iterative template-based picking and 2D-classficiation to obtain a representative set of 2D views yielded 13,922,432 particle coordinates.

After three rounds of iterative 2D classification and selection based on visible image quality and presence of discernible features, 1,934,863 particles were retained. These particles were subsequently classified into 5 classes using *ab initio*, which revealed three maps with interpretable TraI features. The particles from these three *ab initio* classes were combined, amounting to 1,342,170 particles and subjected to homogeneous refinement. This resulted in an initial map of TraI with high resolution features visible in its helicase domain, which was implemented for further processing. To further improve the molecular resolution of the helicase core, the region corresponding to the transesterase and the CTD domains was removed by masking and subsequent signal subtraction in CryoSPARC. These resultant particles were subjected to focused refinement, resulting in a map of the TraI helicase core (TraIHEL-FOC) in complex with the *oriT* DNA substrate, at a resolution of 3.10 Å. This map was sharpened using a B-factor of −100 (Figure S2).

To resolve the variable regions in the initial map within the TE and CTD, the set of 1,342,170 particles were subjected to 3D classification into three classes. Class 1 showed a rigid TE conformation, assigned as state-1, while Classes 2 and 3 contained a flexible TE domain. Therefore, classes 2 and 3 were combined into one particle stack, assigned as state-2. Particles corresponding to these two states each were further subjected to a 3D classification into five classes and subsequently refined by heterogeneous refinement. The best class was selected from the two sets of obtained maps, based on attained map resolution and the presence of variable features, which are missing in other maps due to flexibility. The selected classes were subjected to a final non-uniform refinement job. The two final maps which were obtained, corresponded to the TraIHEL state, in which the TE domain was rigid, and the TraIAHD state, in which the TE domain was flexible. TraIHEL attained a resolution of 3.50 Å and was composed of 157,240 particles, while the TraIAHD attained a resolution of 3.57 Å and was composed of 182,396 particles. These final maps were subjected to map sharpening in CryoSPARC, implementing a B-factor value of −100. For the TraI^TE*^:TraI^310-1756^:fDNA65T/26R complex dataset, CTF parameters were estimated with Patch CTF and based on ice thickness, CTF-fit and astigmatism, 13,073 micrographs were retained for further processing. An initial set of particles were obtained from a subset of 500 micrographs by blob picking using the circular and elliptical blob function, covering a particle diameter between 90 and 120 Å. Templates were selected based on image quality and the presence of discernible features. A workflow of iterative template-based picking and 2D classification yielded a final stack of 5,774,058 particle coordinates from the full set of micrographs. After two rounds of 2D classification and selection of particles, a set of 1,901,410 particles were obtained (Figure S2).

This particle set was subjected to 3 class *ab initio* reconstruction, from which class-3 showed interpretable TraI features. Class-3, containing 1,205,923 particles, was refined using non-uniform refinement and subjected to masked 3D classification into five classes, with the mask encompassing the TE domain and the CTD together. Class-3, composed of 244,018 particles, showed a rigid TE domain. This class was subjected to a further round of particle sorting by 3D classification to improve the resolution of the density encompassing the rigid TE domain and the CTD. This class was finally subjected to non-uniform refinement, producing a map of 3.54 Å resolution and totalling 150,231 particles. Classes 4 and 5, totalling 457,449 particles, displayed a flexible TE domain. These particles were combined and subjected to non-uniform refinement, followed by 3D classification into three classes. Class 1 was composed of 150,319 particles and attained a resolution of 3.49 Å. It showed density corresponding to the flexible TE connected to the helicase domains of TraI via ssDNA, indicating the presence of the asymmetric homodimer of TraI. This map was subjected to sharpening, with a B-factor value of −100 (Figure S2).

In the case of the TraI^310-1756^: fDNA47T/26R complex, CTF parameters were estimated with Patch CTF tool. An initial set of particles was obtained by blob picking from a subset of 500 micrographs, with the particle diameter parameters ranging between 90 and 150 Å and considering up to 1,500 local maxima per micrograph. The particles were refined through iterative template-based picking and 2D classification rounds, implementing a particle diameter of 100 Å, and considering up to 1,500 local maxima per micrograph. After several rounds of cleanup using 2D classification, a total of 1,925,365 particles were obtained. The particles were subjected to 3 class *ab initio* reconstruction, from which Class-3 showed interpretable TraI-helicase features. This particle stack was refined using non-uniform refinement to obtain the final map, containing 690,735 particles and attaining a resolution of 3.30 Å.

The cryo-EM data processing software implemented here are compiled and supported by the SBGrid Consortium^34^. The data processing statistics of all data described in this manuscript can be found in Table S3.

### Model building and refinement

The initial model of the full-length helicase was generated using AlphaFold3^35^ and then manually truncated removing residues beyond Ile1628, as interpretable density could not be seen beyond this. Refinement was initiated by focusing on the helicase core alone, using the higher-resolution TraIHEL-FOC map (generated to exclude the TE domain and CTD), thereby minimising model bias from lower-resolution peripheral regions. The helicase core was subjected to iterative real-space refinement using Phenix^36^ and progress in refinement was tracked using Ramachandran plot and Molprobity^37^. The resulting refined helicase-core model was subsequently rigid-body fitted into the TraIHEL maps and further improved through additional rounds of real-space refinement using Phenix. The helicase-core refined against the TraIHEL-FOC map together with CTD was then used as a starting point for rebuilding and refinement of the extended construct, by iterative real-space refinement using Phenix and progress in refinement was tracked using Ramachandran plot and Molprobity. To model the TE domain bound to *TE binding site*, PDB 2A0I was placed into the density previously assigned to the TE domain of Mol-2 and adjusted as needed to maximise density correspondence. The sequences were modified to match the R1 sequence then finally, the ssDNA trajectories were completed by connecting the ssDNA emerging from the helicase core (Mol-2) to that associated with the TE domain (Mol-1), producing the final TraIAHD model. The PDB coordinate file of the TraIHEL and TraIAHD were deposited to the protein data bank with entry codes 28TI and 28TH respectively.

### Purification of sc-*poriT* template used in the study

The sc-p*oriT* plasmid was transformed into *E. coli* Top10 cells (Thermo Fisher Scientific) and grown in LB medium supplemented with 100 µg/ml ampicillin. Cells were harvested by centrifugation at 6,000 × g for 10 min at 4 °C. Plasmid DNA was extracted using a Plasmid Mega Kit (Qiagen) according to the manufacturer’s instructions, adapted further by undertaking purification steps at 4 °C^38^. DNA was ethanol-precipitated, resuspended in plasmid buffer (20 mM Tris-HCl pH 8.0, 100 mM NaCl, 0.1 mM EDTA), and plasmid supercoiling was analysed by agarose gel electrophoresis on a 1.5% TAE agarose gel.

### Mild trypsin proteolysis of the various complexes described in the study

In the case of the assay probing helicase loading on supercoiled plasmid *oriT* (*sc-poriT*), 0.1 µM *sc-poriT* was incubated with 5 µM, a 50-fold excess, of TraI^TE^, or wild-type TraI^TE^ constructs in 50 mM Tris-HCl pH 7.5, 100 mM NaCl, 10 mM MgCl2 at 37 °C. 1 µM TraI^FL^ or TraI^310-1756^ was then added and incubated for a further 30 min at 37 °C. In the assay including helicase binding site forked DNAs, oligonucleotides of the various combinations of T- and R-strands (Figure 5C), were annealed in an identical workflow to the cryo-EM DNA substrates, and assembled with TraI^FL^ in a 1.5:1, DNA:protein ratio to a final protein concentration of 1 mg/ml and incubated for 10 min at room temperature.

In order to probe the conformation of TraI on the various helicase binding site forked DNA templates, the complexes were treated with bovine trypsin (Sigma) at a TraI:trypsin (w/w) ratio of 400:1 (plasmid DNA reactions) or 100:1 (fDNA reactions) and incubated at room temperature, with sample taken at 30, 60, and 120 min timepoints for analysis. These samples were quenched with 4× SDS-PAGE loading buffer, heated at 95 °C for 10 min, and analysed by SDS-PAGE alongside undigested controls. In the case of the experimental conditions involving *sc-poriT*, the remaining reaction sample at the final 120 min timepoint was stopped by adding proteinase K to 0.1 mg/ml (w/v) and SDS to 0.1% (w/v), followed by incubation for 30 min at room temperature. Plasmid DNA products were analysed by agarose gel electrophoresis on a 1.5% TAE agarose gel.

### Conjugation/Mobilization assay

To assess the conjugation efficiency of different *traI* mutants, we first constructed a *ΔtraI* deletion in the R1 conjugative plasmid using a λ-Red recombineering approach^39^. Briefly, a chloramphenicol-resistance cassette flanked by FRT sites was PCR-amplified using primers containing 50-bp homology arms corresponding to the regions immediately upstream and downstream of *traI* from R1 plasmid. The PCR product was electroporated into *E. coli* Top10 cells (Invitrogen) carrying the R1 plasmid and the pWRG99 plasmid, which had been induced with 0.1 M L-arabinose for 1 h to express the λ-Red recombinase. Recombinants were selected on chloramphenicol plates and transformed with pCP20, enabling FLP-mediated excision of the chloramphenicol cassette. Successful deletion of *traI* and removal of chloramphenicol cassette was confirmed by sequencing. *E. coli* Top10 cells harbouring R1Δ*traI* were subsequently transformed with pCDF plasmids encoding either wild type *traI* or selected *traI* mutant alleles, constructed as described above.

Conjugation assays were performed as previously reported with minor modifications^40^. Donor strains consisted of *E. coli* Top10 carrying either R1 or R1Δ*traI* (kanamycin-resistant), with or without the complementing pCDF plasmids. The recipient strain was *E. coli* Top10 carrying the ampicillin-resistant pFPV plasmid expressing mCherry. Overnight cultures of donors (200 μl) and recipients (2 ml) were diluted into 5 ml and 50 ml fresh LB, respectively, without antibiotics and grown at 37 °C with shaking (180 rpm) to OD₆₀₀ = 0.3–0.4. For induction of the complementing plasmids, donors were treated with 1 mM IPTG and grown for an additional hour before cultures were placed on ice. To remove residual antibiotics, 1 ml of each donor culture was centrifuged (2 min, 4,000 × g, 4 °C) in a microcentrifuge tube and the pellet was gently resuspended in 1 ml chilled LB. For recipients, 50 ml culture was centrifuged (10 min, 4,000 × g, 4 °C) and the pellet was resuspended in 5 ml chilled LB. This washing step was repeated twice for all strains. Washed cells were adjusted to OD₆₀₀ = 1.0 for donors and OD₆₀₀ = 5.0 for recipients. Conjugation mixtures were prepared by combining 100 μl donor with 100 μl recipient, briefly vortexing, and incubating at 37 °C with shaking (180 rpm) for 30 min. Reactions were then chilled on ice, serially diluted, and plated on LB agar containing carbenicillin and kanamycin to enumerate transconjugants, and on antibiotic-free LB plates to determine total viable cell counts. Experiments were repeated with at least three biological replicates.

### Forked DNA duplex separation assay

A T-strand of 22 or 34 nucleotides was annealed to its partially complementary single-stranded DNA (ssDNA) sequence containing either 11 or 23 complementary bases respectively at a 1:1 molar ratio. The 3′ end of the antisense R-strand oligonucleotide contained a poly-T tail to generate DNA fork substrates (Figure 6A and 6E). Oligonucleotides were labelled at their 3′ ends with fluorescent dyes (Cy5, FITC or FAM), except the 34mer, which was labelled on its 5’ end. Each ssDNA oligonucleotide was dissolved to a concentration of 100 µM in 20 mM Tris-HCl pH 7.4, 100 mM NaCl. The T- and R-strand oligonucleotides (Figure 6A and 6E) were mixed to a final concentration of 10 µM in the same buffer, heated to 95 °C for 5 min, and gradually cooled to room temperature over 30 min to allow for their annealing. Single-stranded oligonucleotides were similarly heated to 95 °C for 5 min but were immediately placed on ice to preserve their single-stranded state. The resulting annealed DNA duplexes and single-stranded oligonucleotides were subsequently diluted to 1 µM to prepare the working stock solutions.

Reaction mixtures attained a final volume of 30 µL, containing 0.33 µM of the annealed DNA substrate in 20 mM Tris-HCl pH 7.4, 100 mM NaCl, 5 mM MgCl₂. Reactions were assembled at room temperature and initiated by the addition of TraI^FL^ at final concentrations that correspond to 0.2×, 0.5×, 1×, 2× and 4× excesses over the DNA substrate. The mixtures were incubated at room temperature for 20 min. For conditions which include ATP, 2 mM ATP was added after a 10 min pre-incubation of TraI and DNA substrates, followed by a further incubation at room temperature for 10 min. Reaction products were resolved on a 4% TBE agarose gel and visualized using the Invitrogen iBright Imaging System (Thermo Fisher Scientific) under fluorescent imaging condition FAM/FITC:495 nm and 515 nm; Cy5: 649 nm and 670 nm, excitation and emission wavelengths respectively.

### Nicking inhibition assay

Reaction mixtures contained 50 pM *sc-poriT* DNA, 20 mM Tris-HCl pH 8.0, 70 mM NaCl, 10 mM MgCl2, and where indicated, 2.5 µM TraI^TE^ with 25 pM–1 µM IHF, corresponding to *sc-poriT*:TraI^TE^ and *sc-poriT*:IHF ratios of 1:50 and 1:0.5–20, respectively. Reactions were assembled at room temperature, plasmid nicking was initiated by addition of wildtype TraI^TE^, followed by incubation at 37 °C for 30 min. Reactions were stopped by adding proteinase K to 0.1 mg/ml (w/v) and SDS to 0.1% (w/v), followed by incubation for 30 min at room temperature. DNA products were analysed by agarose gel electrophoresis on a 1.5% TAE agarose gel.

## Data availability

The cryo-EM maps and the corresponding atomic coordinates have been deposited to the Electron Microscopy data bank (EMDB) and Protein data bank (PDB) databases. The EMDB codes are EMD-56432, EMD-56434, EMD-56435, EMD-56436, EMD-56437, and the PDB codes are 28TI and 28TH.

## Author contributions

DG performed the biochemical and structural work of the manuscript. RL performed the TraI mediated strand separation assay and cryo-EM data processing of the TraI^310-1756^:fDNA47T/26R complex. JBP performed the *in vivo* analysis reported. SBL contributed to the project at the initial stage with preliminary analysis leading to the final work reported here. TRDC supervised the *in vivo* work, AI supervised the cryo-EM structural characterisation and biochemical work described. The manuscript was written by TRDC and AI with contributions from DG, RL, JBP and SBL.

## Acknowledgements

We would like to thank Dr. Hui Zhang for cryo-EM in the QMUL cryo-EM facility for in-house grid screening; Dr. Charlotte Millership, the protein facility manager for the support with facility; Dr. Vitaly Voloshin for the support with the cryo-EM cluster and Dr. Nora Cronin, the facility manager of the London Consortium for Cryo-EM (LonCEM) for the help with cryo-EM data collection. We acknowledge Diamond Light Source for access and support of the cryo-EM facilities at the UK National Electron Bio-imaging Centre (eBIC, proposal BI42268). The Jeol 2100 plus TEM used for cryo-EM grid screening at QMUL was funded by an UKRI Altert16 grant (BB/R000514/1). This work was supported by an AMS Springboard award (SBF007\100161) and by a BBSRC project grant (BB/X016900/1).

