## Supplementary Figures and Table for "Relaxase Asymmetry Drives Initiation of Bacterial Conjugation"

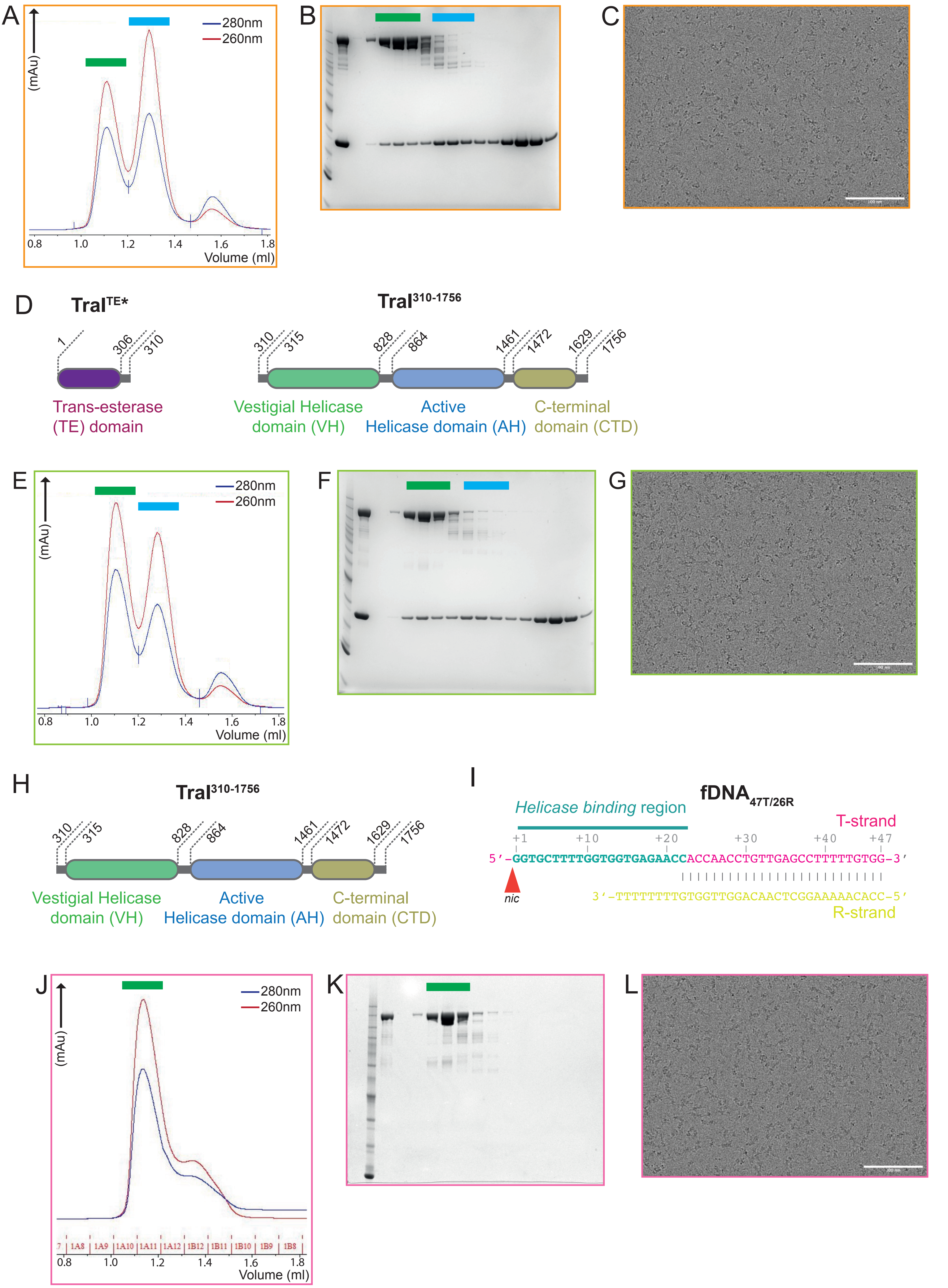
A
280nm
260nm
(mAu)
0.8 1.0 1.2 1.4 1.6 1.8
Volume (ml)
B
C
D
TraI$^{TE*}$
1 306 310
Trans-esterase (TE) domain
TraI$^{310-1756}$
310 315 828 864 1461 1472 1629 1756
Vestigial Helicase domain (VH)
Active Helicase domain (AH)
C-terminal domain (CTD)
E
280nm
260nm
(mAu)
0.8 1.0 1.2 1.4 1.6 1.8
Volume (ml)
F
G
H
TraI$^{310-1756}$
310 315 828 864 1461 1472 1629 1756
Vestigial Helicase domain (VH)
Active Helicase domain (AH)
C-terminal domain (CTD)
I
fDNA$_{47T/26R}$
Helicase binding region
T-strand
+1 +10 +20 +30 +40 +47
5'-GGTGCTTTTGGTGGTGAGAACCACCAACCTGTTGAGCCTTTTTGTGG-3'
nic
3'-TTTTTTTTGTGGTTGGACAACTCGGAAAAACACC-5'
R-strand
J
280nm
260nm
(mAu)
7 1A8 1A9 1A10 1A11 1A12 1B12 1B11 1B10 1B9 1B8
0.8 1.0 1.2 1.4 1.6 1.8
Volume (ml)
K
L

**Supplementary Figure 1: Biochemistry and cryo-EM**

a. Size-exclusion column chromatogram of the tripartite TraITE*–TraIFL*–fDNA$_{65T/26R}$ assembly. The peak marked with a green bar correspond to the higher molecular weight TraITE*:TraIFL*:fDNA$_{65T/26R}$ complex and the peak marked with a blue bar correspond to the lower molecular weight TraITE*:fDNA$_{65T/26R}$ complex.
b. SDS-PAGE of the size-exclusion chromatogram from panel S1-A.
c. Representative cryo-EM micrograph of the TraITE*:TraIFL*:fDNA$_{65T/26R}$ complex.
d. Left - Domain organisation of the TraI$^{TE}$* construct used for the biochemical assembly, with domain boundaries. Right - Domain organisation of the TraI$^{310-1756}$ construct used for the biochemical assembly, with domain boundaries.
e. Size-exclusion column chromatogram of the tripartite TraI$^{TE}$*–TraI$^{310-1756}$–fDNA$_{65T/26R}$ assembly. The peak marked with a green bar correspond to the higher molecular weight TraI$^{TE}$*:TraI$^{310-1756}$:fDNA$_{65T/26R}$ complex and the peak marked with a blue bar correspond to the lower molecular weight TraI$^{TE}$*:fDNA$_{65T/26R}$ complex.
f. SDS-PAGE of the size-exclusion chromatogram from panel S1-E.
g. Representative cryo-EM micrograph of the TraI$^{TE}$*:TraI$^{FL}$*:fDNA$_{65T/26R}$ complex.
h. Domain organisation of the TraI$^{310-1756}$ construct used for the biochemical assembly, with domain boundaries.
i. Sequence of the DNA fork substrate (fDNA$_{47T/26R}$) used for the assembly of the bipartite TraI$^{310-1756}$:fDNA$_{47T/26R}$ complex.
j. Size-exclusion column chromatogram of the bipartite TraI$^{310-1756}$–fDNA$_{47T/26R}$ assembly. The peak marked with a green bar correspond to the higher molecular weight TraI$^{310-1756}$:fDNA$_{47T/26R}$ complex.
k. SDS-PAGE of the size-exclusion chromatogram from panel S1-J.
l. Representative cryo-EM micrograph of the TraI$^{310-1756}$:fDNA$_{47T/26R}$ complex.

### Cryo-EM workflow of TraI$^{TE*}$:Tra$^{FL*}$:fDNA$_{65T/26R}$ complex

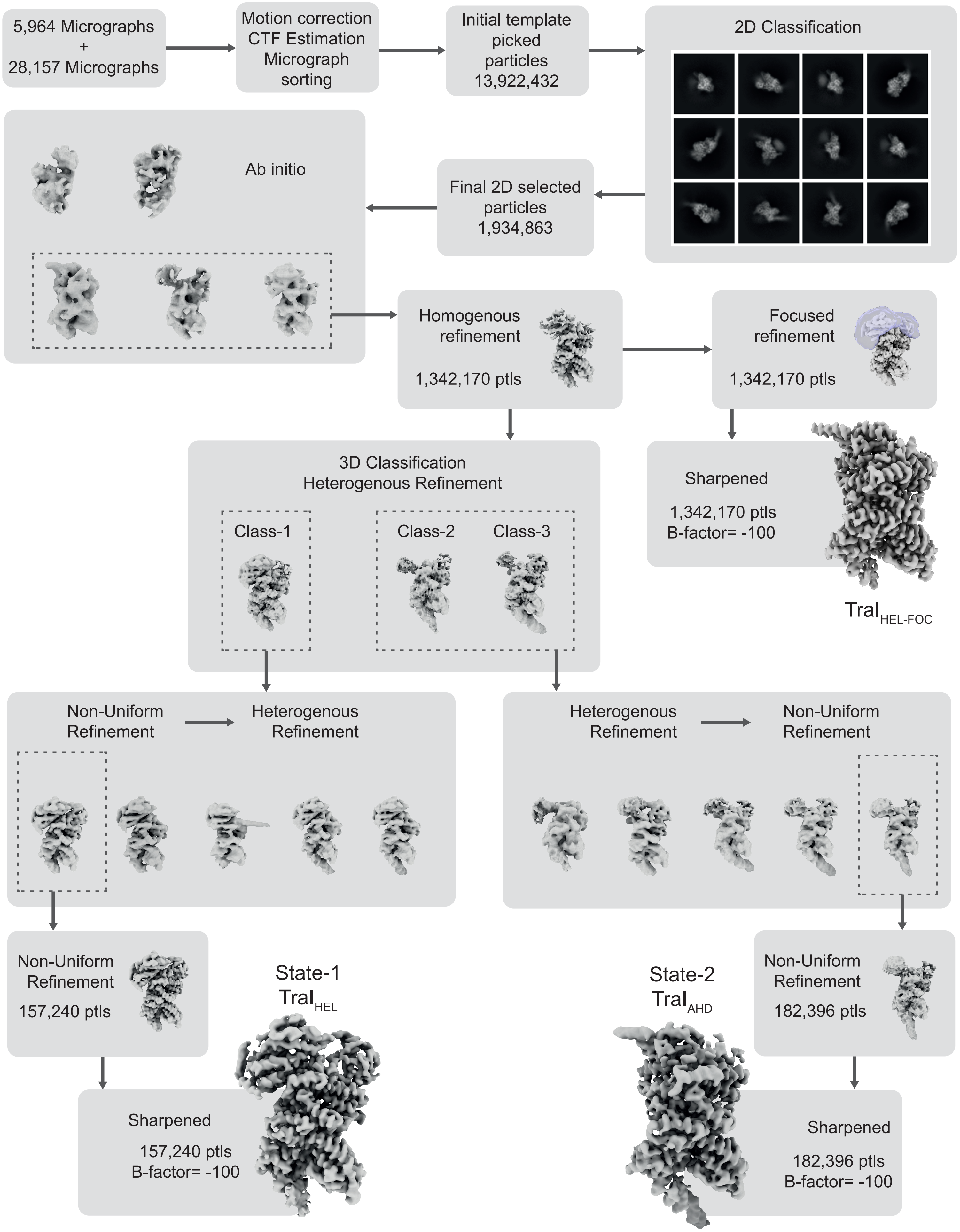


**Supplementary Figure 2: Cryo-EM workflow of TraI$^{TE*}$:Tra$^{FL*}$:fDNA$_{65T/26R}$ complex**

A

**Cryo-EM workflow of $TraI^{TE}$:$TraI^{310\text{-}1756}$:$fDNA_{65T/26R}$**

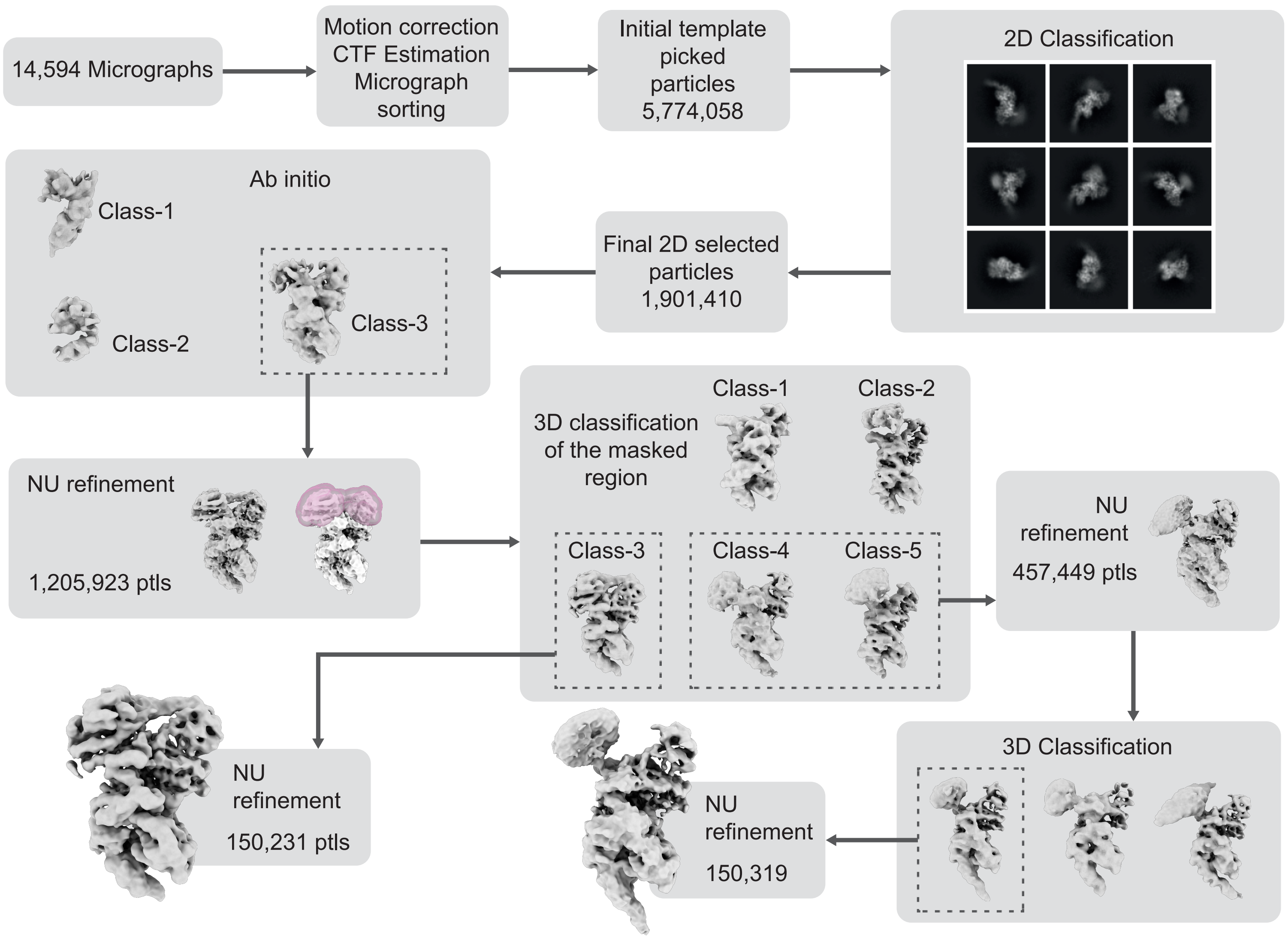


B

**Cryo-EM workflow of $TraI^{310\text{-}1756}$:$fDNA_{47T/26R}$**

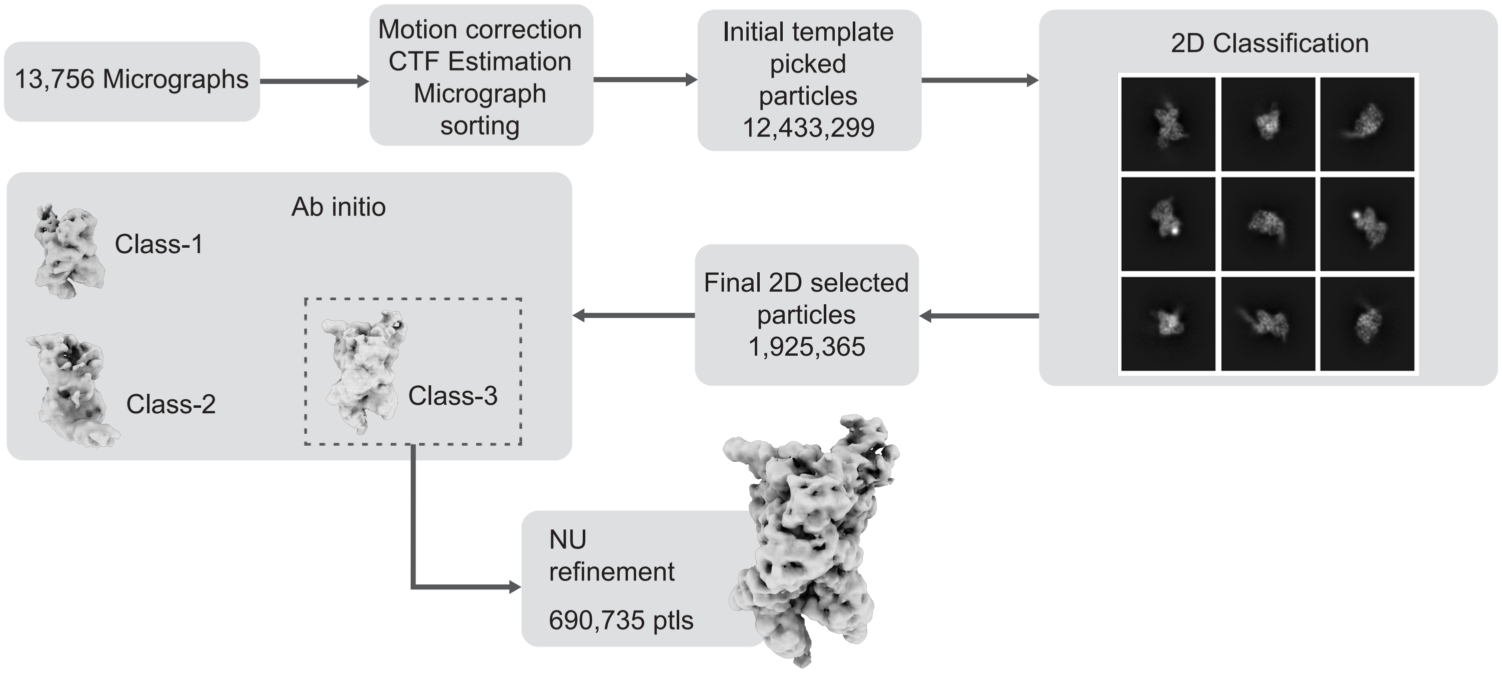


**Supplementary Figure 3: Cryo-EM workflow the of the control TraI-fDNA complexes**

a. Cryo-EM workflow of $TraI^{TE*}$:$TraI^{310\text{-}1756}$:$fDNA_{65T/26R}$ complex.
b. Cryo-EM workflow of $TraI^{310\text{-}1756}$:$fDNA_{47T/26R}$ complex.

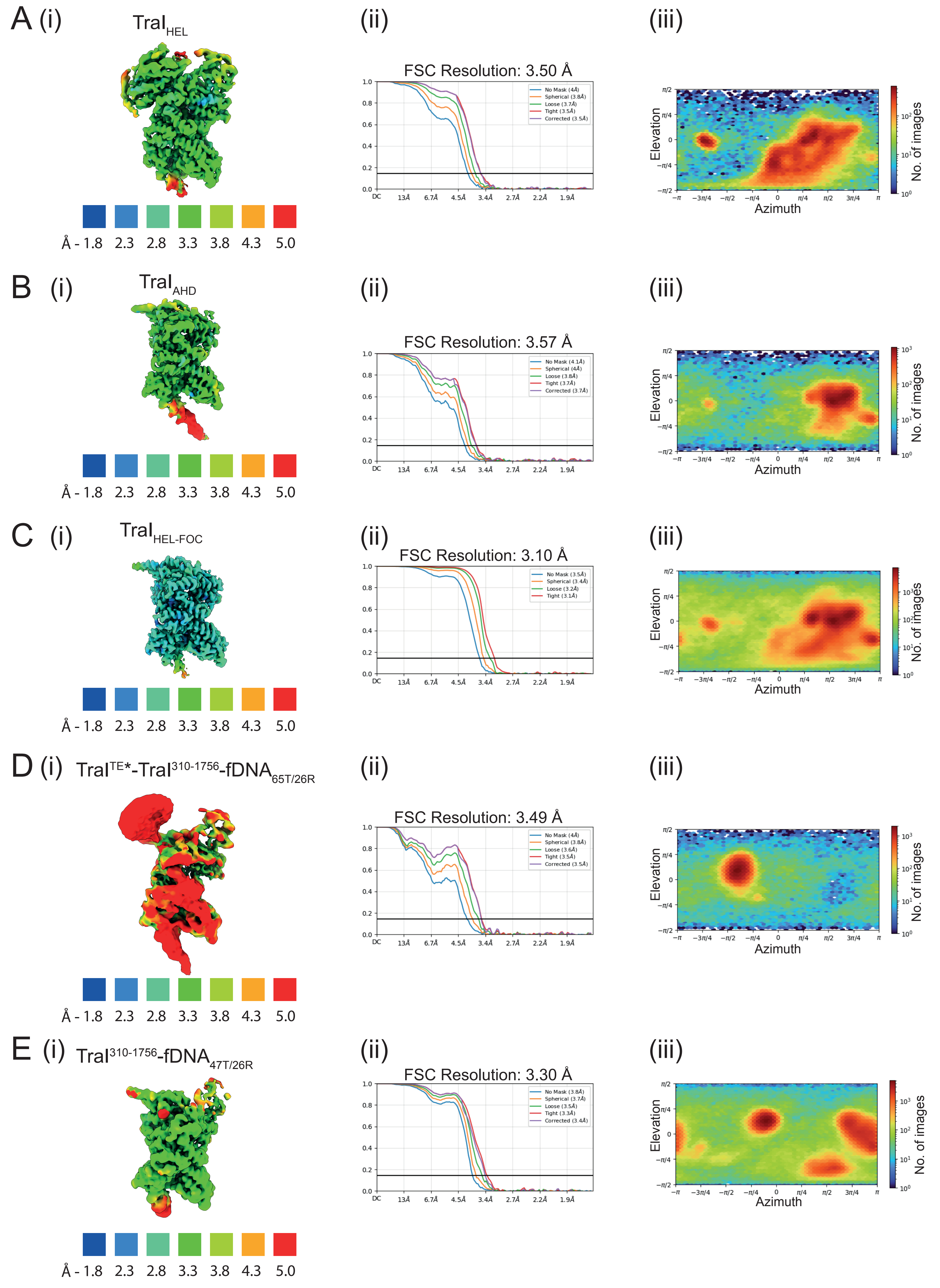

A (i) $TraI_{HEL}$
Å - 1.8 2.3 2.8 3.3 3.8 4.3 5.0
(ii)
FSC Resolution: 3.50 Å
No Mask (4Å)
Spherical (3.8Å)
Loose (3.7Å)
Tight (3.5Å)
Corrected (3.5Å)
DC 13Å 6.7Å 4.5Å 3.4Å 2.7Å 2.2Å 1.9Å
(iii)
Elevation
Azimuth
No. of images
B (i) $TraI_{AHD}$
Å - 1.8 2.3 2.8 3.3 3.8 4.3 5.0
(ii)
FSC Resolution: 3.57 Å
No Mask (4.1Å)
Spherical (4Å)
Loose (3.8Å)
Tight (3.7Å)
Corrected (3.7Å)
(iii)
Elevation
Azimuth
No. of images
C (i) $TraI_{HEL-FOC}$
Å - 1.8 2.3 2.8 3.3 3.8 4.3 5.0
(ii)
FSC Resolution: 3.10 Å
No Mask (3.5Å)
Spherical (3.4Å)
Loose (3.2Å)
Tight (3.1Å)
(iii)
Elevation
Azimuth
No. of images
D (i) $TraI^{TE*}$-$TraI^{310-1756}$-$fDNA_{65T/26R}$
Å - 1.8 2.3 2.8 3.3 3.8 4.3 5.0
(ii)
FSC Resolution: 3.49 Å
No Mask (4Å)
Spherical (3.8Å)
Loose (3.6Å)
Tight (3.5Å)
Corrected (3.5Å)
(iii)
Elevation
Azimuth
No. of images
E (i) $TraI^{310-1756}$-$fDNA_{47T/26R}$
Å - 1.8 2.3 2.8 3.3 3.8 4.3 5.0
(ii)
FSC Resolution: 3.30 Å
No Mask (3.8Å)
Spherical (3.7Å)
Loose (3.5Å)
Tight (3.3Å)
Corrected (3.4Å)
(iii)
Elevation
Azimuth
No. of images

**Supplementary Figure 4: Cryo-EM maps described in this study**

a. Cryo-EM map of the state-1, $TraI_{HEL}$ complex. (i) Map coloured by local resolution contoured at 0.25 σ. (ii) Fourier shell correlation of the cryo-EM map showing an average map resolution of 3.50 Å. (iii) Orientation distribution of particles contributing to the cryo-EM map.
b. Cryo-EM map of the state-2, $TraI_{AHD}$ complex (i) Map coloured by local resolution contoured at 0.18 σ. (ii) Fourier shell correlation of the cryo-EM map showing an average map resolution of 3.57 Å. (iii) Orientation distribution of particles contributing to the cryo-EM map.
c. Focused refined cryo-EM map of the $TraI_{HEL\text{-}FOC}$ region (i) Map coloured by local resolution contoured at 0.30 σ. (ii) Fourier shell correlation of the cryo-EM map showing an average map resolution of 3.10 Å. (iii) Orientation distribution of particles contributing to the cryo-EM map.
d. Cryo-EM map of the $TraI^{TE*}$:$TraI^{310\text{-}1756}$:$fDNA_{65T/26R}$ complex (i) Map coloured by local resolution contoured at 0.14 σ. (ii) Fourier shell correlation of the cryo-EM map showing an average map resolution of 3.49 Å. (iii) Orientation distribution of particles contributing to the cryo-EM map.
e. Cryo-EM map of the $TraI^{310\text{-}1756}$:$fDNA_{47T/26R}$ complex (i) Map coloured by local resolution contoured at 0.19 σ. (ii) Fourier shell correlation of the cryo-EM map showing an average map resolution of 3.30 Å. (iii) Orientation distribution of particles contributing for the cryo-EM map.

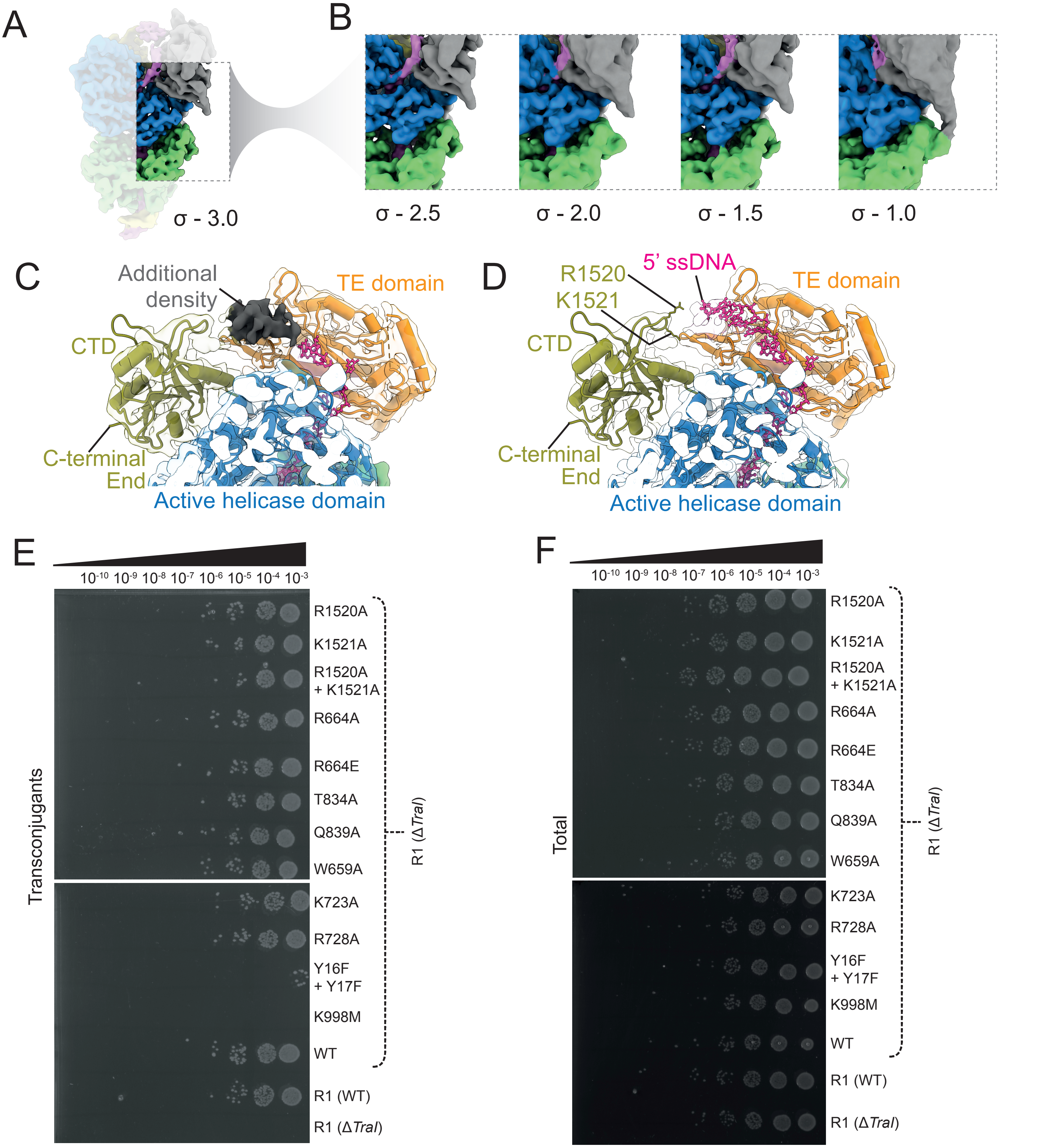


**Supplementary Figure 5: Structure of TraI$_{HEL}$ and in vivo validation**

a. The cryo-EM map of TraI$_{HEL}$ at σ 3.0, coloured by domains, highlighting the region between the TE and VH domains.
b. Cryo-EM maps of TraI$_{HEL}$ at various σ levels focused at the region between TE and VH domains.
c. The TraI$_{HEL}$ model (shown in cartoon) overlayed on cryo-EM map (transparent). The individual domains are coloured. The unassigned additional density is shown in dark grey.
d. The TraI$_{HEL}$ model (shown in cartoon) overlayed on cryo-EM map (transparent). The individual domains are coloured. ssDNA built into the previously unassigned additional density shown in Figure S5-C.
e-f. Results of the *in vivo* conjugation assay of TraI point mutants at its DNA interaction interface. *E. coli* cells carrying a kanamycin-resistant wild-type R1 plasmid or an R1 ΔtraI plasmid complemented with plasmids encoding TraI variants bearing the indicated point mutations were mated with carbenicillin-resistant recipient *E. coli* cells. After 30 min, serial ten-fold dilutions of mating mixtures were plated on double-selection LB agar plate to enumerate transconjugants (E). In parallel, identical dilutions were plated on a non-selective plate to determine total viable cell counts for standardisation of the results (F). Brackets indicate TraI variants expressed in the R1 ΔtraI background; wild-type R1 and control knockout ΔtraI strains are shown at the bottom of each panel; mutations Y16F+Y17F and K998M are catalytically defective and were used as negative controls.

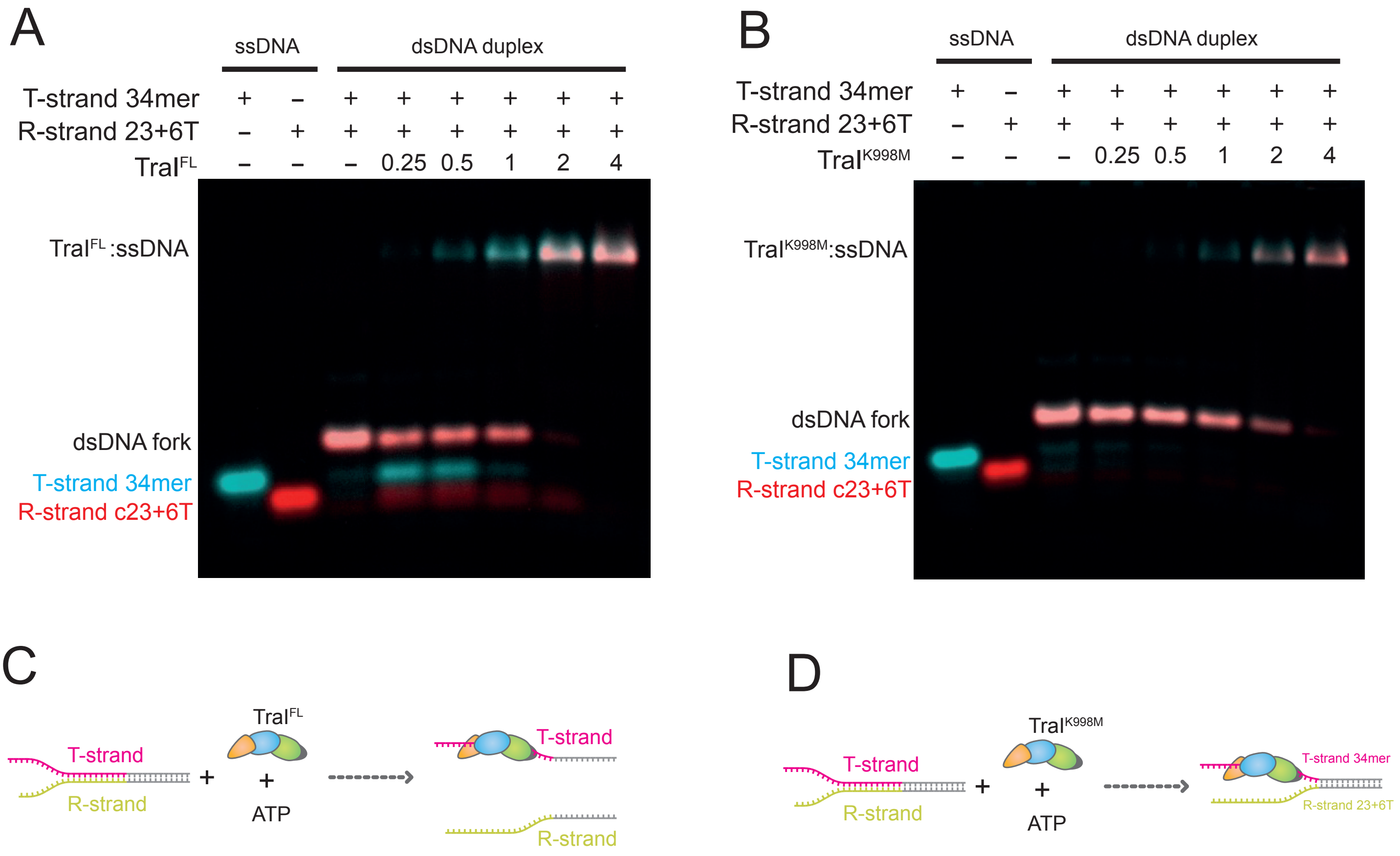


**Supplementary Figure 6: ATP dependent unwinding of DNA substrate by TraI helicase**

a. Gel shift assay of TraI with 34mer (T-strand), 29mer (R-strand) or dsDNA duplex of $fDNA_{34T/23R}$ with ATP at stoichiometric $TraI^{FL}$ concentration of 0.25, 0.5, 1, 2 or 4 fold excess compared to the DNA substrate. Details of this assay are available in Methods.

b. Gel shift assay of TraI with 34mer (T-strand), 29mer (R-strand) or dsDNA duplex of fDNA34T/23R with ATP at stoichiometric $TraI^{K998M}$ concentration of 0.25, 0.5, 1, 2 or 4 fold excess compared to DNA substrate. Details of this assay are available in Methods.

c. Schematic summary of $TraI^{FL}$ unwinding $fDNA_{34T/23R}$ template in the presence of ATP.

d. Schematic summary of $TraI^{K998M}$ unable to unwind $fDNA_{34T/23R}$ template in the presence of ATP.

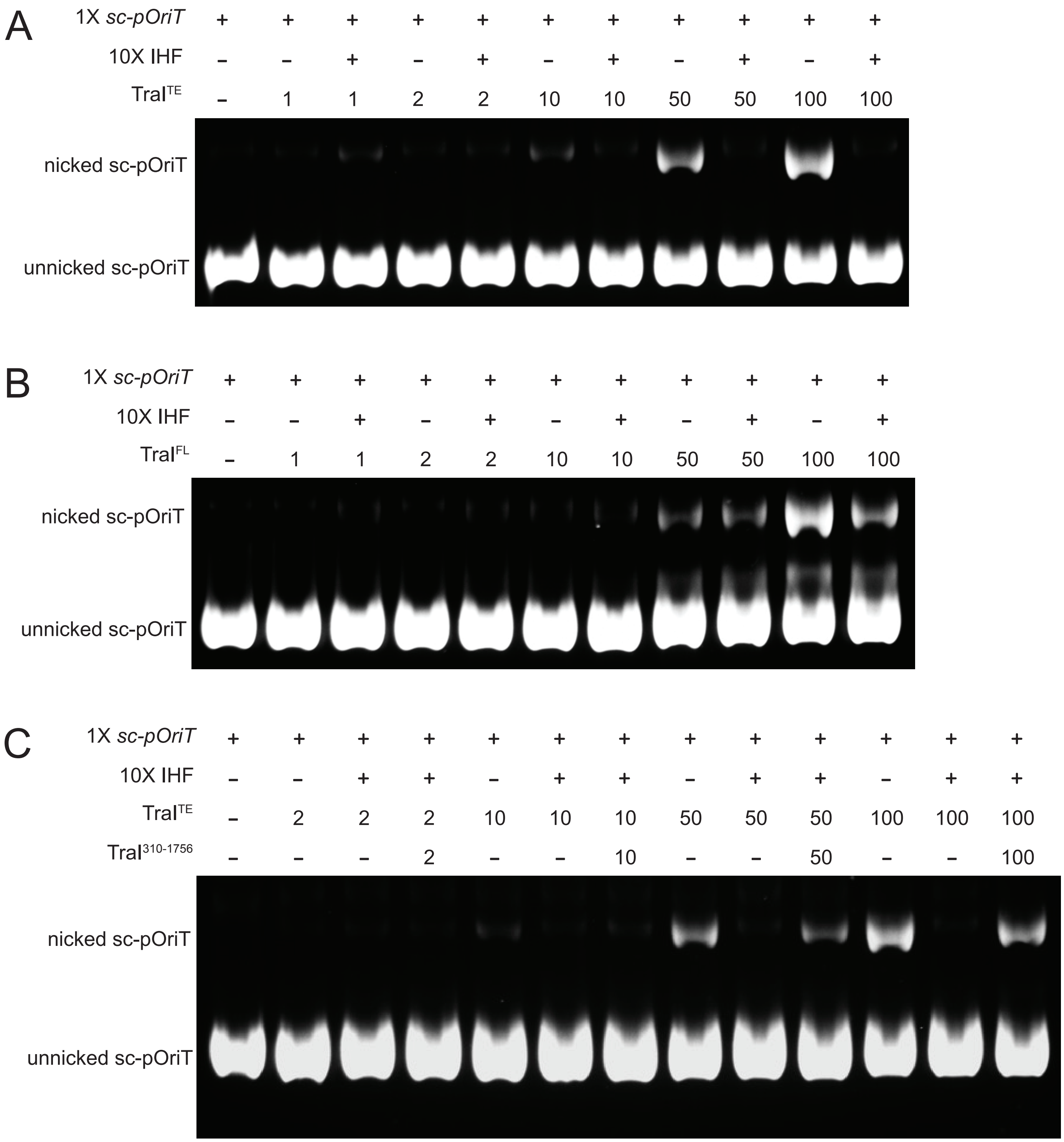


**Supplementary Figure 7: TraITE mediated nicking assay**

a. Agarose gel showing $TraI^{TE}$ mediated nicking of *scporiT* at various concentration and in the presence or absence of IHF.
b. Agarose gel showing $TraI^{FL}$ mediated nicking of *scporiT* at various concentration and in the presence or absence of IHF.
c. Agarose gel showing $TraI^{TE}$ mediated nicking of *scporiT* at various concentration and in the presence or absence of IHF and/or $TraI^{310-1756}$.

**Supplementary Table 1:** List of plasmids implemented in biochemical experiments

| Biochemistry Constructs | Truncations | Mutations | Tag | Vector | Source |
|---|---|---|---|---|---|
| $TraI^{TE}$ | 1-309 | WT | Strep-tag II | pCDF-1b | This work |
| $TraI^{TE}$ | 1-309 | Y16F, Y17F | Strep-tag II | pCDF-1b | This work |
| $TraI^{310\text{-}1756}$ | 310-1756 | WT | 6xHis | pCDF-1b | This work |
| $TraI^{FL}$ | 1-1756 | WT | 6xHis | pCDF-1b | Ilangovan et al., 2017 |
| $TraI^{FL}$ | 1-1756 | Y16F, Y17F | 6xHis | pCDF-1b | This work |
| $TraI^{FL}$ | 1-1756 | K998M | 6xHis | pCDF-1b | This work |
| IHF | FL | WT | 6xHis | pRSF-duet | This work |
| *scpOriT* | n/a | n/a | n/a | pUC19 | This work |

**Supplementary Table 2:** List of plasmids implemented in conjugation experiments

| Conjugation Constructs | Truncations | Mutations | Vector | Source |
|---|---|---|---|---|
| R1 (WT) | n/a | n/a | n/a | Ilangovan et al., 2017 |
| R1 (ΔTraI) | n/a | n/a | n/a | This work |
| TraI | FL | WT | pCDF-1b | This work |
| TraI | FL | Y16F+Y17F | pCDF-1b | This work |
| TraI | FL | W659A | pCDF-1b | This work |
| TraI | FL | R664A | pCDF-1b | This work |
| TraI | FL | R664E | pCDF-1b | This work |
| TraI | FL | K723A | pCDF-1b | This work |
| TraI | FL | R728A | pCDF-1b | This work |
| TraI | FL | T834A | pCDF-1b | This work |
| TraI | FL | Q839A | pCDF-1b | This work |
| TraI | FL | K998M | pCDF-1b | This work |
| TraI | FL | R1520A | pCDF-1b | This work |
| TraI | FL | K1521A | pCDF-1b | This work |
| TraI | FL | R1520A+K1621A | pCDF-1b | This work |

**Supplementary Table 3:** Cryo-EM data collection, processing, and model refinement

| **Data collection and processing** | | | | | |
|---|---|---|---|---|---|
| **Microscope** | Titan Krios G3i (3.1) | | | | |
| **No. of Movies** | 34,121 | | | 14,594 | 13,765 |
| **Camera** | Gatan K3 Bioquantum | | | Gatan K3 Bioquantum | Gatan K3 Bioquantum |
| **Voltage (kV)** | 300 | | | 300 | 300 |
| **dose ( e−/Å²)** | 63 | | | 63 | 60 |
| **Pixel size (Å)** | 0.84 | | | 0.84 | 0.84 |
| **Defocus range (µm)** | -0.5 to -3.0 | | | -0.5 to -3.0 | -0.5 to -3.0 |
| **Symmetry imposed** | C1 | | | C1 | C1 |
| **Complex** | **$TraI_{HEL}$ (State 1)** | **$TraI_{AHD}$ (State 2)** | **$TraI_{HEL\text{-}FOC}$ (Helicase)** | **$TraI^{TE}$:$TraI^{310\text{-}1756}$:$fDNA_{65T/26R}$** | **$TraI^{310\text{-}1756}$:$fDNA_{47T/26R}$** |
| **Grid** | Quantifoil R2/2 Au 200 | | | Quantifoil R2/2 Au 200 | Quantifoil R2/2 Au 200 |
| **Initial particle images** | 13,922,432 | | | 5,774,058 | 12,433,299 |
| **Final particle images** | 157,240 | 182,396 | 1,342,170 | 150,319 | 690,735 |
| **FSC threshold** | 0.143 | | | 0.143 | 0.143 |
| **Map resolution (Å)** | 3.50 | 3.57 | 3.09 | 3.49 | 3.31 |
| **Map B-factor** | 164.6 | 164.9 | 170.4 | 135.1 | 198.5 |
| **EMDB codes** | EMD-56436 | EMD-56432 | EMD-56437 | EMD-56435 | EMD-56434 |
| **Refinement and model validation** | | | | | |
| **Model vs data CC (mask)** | 0.87 | 0.85 | | | |
| **Clash score** | 7.79 | 6.4 | | | |
| **Molprobity score** | 1.74 | 1.68 | | | |
| **Bond length rmsd** | 0.004 | 0.003 | | | |
| **Bond angle rmsd** | 0.685 | 0.527 | | | |
| **Poor outliers (%)** | 2.56 | 2.7 | | | |
| **Ramachandran** | | | | | |
| **Favoured (%)** | 98.43 | 98.38 | | | |
| **Allowed (%)** | 1.57 | 1.62 | | | |
| **Outlier (%)** | 0 | 0 | | | |
| **PDB code** | 28TI | 28TH | | | |
